# RFX3 is essential for the development and maturation of human pancreatic islets derived from pluripotent stem cells

**DOI:** 10.1101/2024.09.18.613711

**Authors:** Bushra Memon, Noura Aldous, Ahmed K. Elsayed, Sadaf Ijaz, Sikander Hayat, Essam M. Abdelalim

## Abstract

RFX3 in human pancreatic islet development has not been explored. This study aims to investigate the function of RFX3 in human pancreatic islet development using human islet organoids derived from iPSCs, hypothesizing that RFX3 regulates human islet cell differentiation. We generated *RFX3* knockout (*RFX3* KO) iPSC lines using CRISPR/Cas9 and differentiated them into pancreatic islet organoids. Various techniques were employed to assess gene expression, cell markers, apoptosis, proliferation, and glucose-stimulated insulin secretion. Single-cell RNA sequencing (scRNA-seq) datasets from hESC-derived pancreatic islets were re-analyzed to investigate *RFX3* expression in specific cell populations at various developmental stages. Furthermore, bulk RNA sequencing was conducted to further assess transcriptomic changes. RFX3 was found to be highly expressed in pancreatic endocrine cell populations within pancreatic progenitors (PPs), endocrine progenitors (EPs), and mature islet stages derived from iPSCs. scRNA-seq further confirmed RFX3 expression across different endocrine cell clusters during differentiation. *RFX3* loss disrupted pancreatic endocrine gene regulation, reduced hormone-secreting islet cells, and impaired beta-cell function and insulin secretion. Despite a significant reduction in pancreatic islet hormones, the pan-endocrine marker CHGA remained unchanged at both EP and islet stages, likely due to an increase in enterochromaffin cells (ECs). This was supported by our findings of high EC marker expression in *RFX3* KO EPs and islets. In addition, *RFX3* loss led to smaller islet organoids, elevated TXNIP levels, and increased apoptosis in EPs and islets. These findings underscore the crucial role of RFX3 in regulating human islet cell differentiation and its role in suppressing enterochromaffin cell specification. These insights into RFX3 function have implications for understanding islet biology and potential diabetes susceptibility.

## Introduction

The Islets of Langerhans in the endocrine pancreas play an essential role in regulating blood glucose levels, and defects in their development or function are associated with diabetes. Although much of our knowledge about genes involved in pancreatic islet development and diabetes pathogenesis comes from animal studies, significant differences exist between humans and animals, particularly in islet structure and physiological properties of endocrine cells [1]. Furthermore, recapitulating diabetes phenotypes in animal models remains challenging. In this context, human pluripotent stem cells (hPSCs) have emerged as a valuable tool for modeling different types of diseases, including diabetes [2, 3].

Regulatory factor X3 (RFX3) has recently been implicated in pancreatic islet development in mice. It is expressed in neurogenin 3 (Neurog3)-positive endocrine progenitors (EPs) as early as embryonic day 13.5 and continues to be expressed in adult mouse islets along with insulin (Ins), glucagon (Gcg), somatostatin (Sst), and pancreatic polypeptide-y (Ppy) [4, 5]. In *Rfx3^-/-^* mice, surviving embryos exhibit impaired islet cytoarchitecture with reduced numbers of Ins^+^, Gcg^+^, and Sst^+^ cells, increased Ppy^+^ cells, and glucose intolerance. Despite these defects, *Rfx3* deficiency did not impact any pancreatic developmental markers such as Pdx1, Nkx6.1, Neurod1 and Mafa at any developmental stage [4, 5]. Of note, another key RFX family member, RFX6 has been shown to regulate human pancreatic islet development and function [6–8]. Homozygous mutations in *RFX6* cause Mitchell-Riley Syndrome (MRS) that feature severe neonatal diabetes whereas heterozygous mutations in *RFX6* cause maturity onset diabetes of the young (MODY) [9, 10]. However, the role of RFX3 in human pancreatic development remains unclear.

Given the crucial role of RFX3 in rodents in regulating pancreatic islet cell function and development, investigating its function in human pancreatic development is highly warranted. Therefore, in this study, we generated *RFX3* knockout (*RFX3* KO) induced PSC (iPSC) model to investigate the role of RFX3 in human pancreatic islet development. Our findings revealed that the loss of *RFX3* caused significant disruption in the pancreatic endocrine program during islet differentiation and led to increased differentiation into enterochromaffin cells (ECs).

## Materials and Methods

### iPSC maintenance and generation of *RFX3* knockout iPSCs

iPSCs were generated in our lab from healthy individual, and they were fully characterized and maintained in culture as described in our previous report [11]. iPSCs were cultured on 1:80 diluted geltrex using mTeSR Plus medium (Stem Cell Technologies, Canada) supplemented with 1% Penicillin-streptomycin.

A CRISPR/Cas9 technique was used to generate *RFX3* KO iPSC lines. Guide RNA (gRNA) targeting *RFX3* coding region was cloned into a GFP-tagged vector expressing spCas9 and transfected into undifferentiated iPSCs, dissociated into single cells, using Lipofectamine 3000 reagent (ThermoFisher Scientific, Massachusetts, USA) using manufacturer’s protocol. The transfected iPSCs were then plated on 1:80 geltrex for 48 hours prior to sorting GFP-expressing iPSCs. The isolated single cells were then cultured and expanded as clones. DNA was extracted using QuickExtract DNA reagent (Invitrogen, Massachusetts, USA) and the target region was amplified using primers flanking the gRNA binding site. PCR products were used for sanger sequencing for KO verification. Four *RFX3* KO clones were generated and loss of RFX3 protein expression was confirmed using immunofluorescence upon differentiating to pancreatic progenitors (PPs). Finally, two *RFX3* KO clones were selected for further experiments (*RFX3* KO1 and *RFX3* KO2 iPSCs).

### Differentiation of iPSCs into pancreatic islet organoids

Differentiation was started when iPSCs plated on 1:50 geltrex reached 70-80% confluency. Differentiation was performed in adherent culture until the PPs using our optimized protocol [12]. Further differentiation to beta cells was performed either in 2D format or 3D where PPs were dissociated into single cells and 2 × 10^6^ WT and *RFX3* KO PPs were re-aggregated in Aggrewell 400 plates for organoid formation using EP day 1 differentiation media (Supplementary Fig. 1A), adapted from Veres et al. protocol for further differentiation into pancreatic islet cells [13]. Organoids were transferred from the microwells into suspension culture in ultra-low attachment plates after 4 days in EP stage and differentiated into mature islets. Media composition for each of differentiation is provided in Supplementary Table 1.

### RNA extraction and real-time quantitative polymerase chain reaction (RT-qPCR)

Differentiated cells were treated with TRIzol reagent (Life Technologies) prior to phase separation using chloroform. RNA extraction was performed on the aqueous phase using Direct-zol RNA Miniprep kit (Zymo Research Corporation, California, USA). 1 µg of total RNA was used for cDNA synthesis using High-Capacity cDNA Reverse Transcription Kit while following manufacturer’s protocol (Applied Biosystems, Massachusetts, USA). RT-qPCR was performed for the synthesized cDNA using GoTaq qPCR SYBR Green Master Mix (Promega) for specific primers provided in Supplementary Table 2. GAPDH was used as an endogenous or housekeeping control while fold changes for KO samples were calculated using the ddCt method taking WT levels as reference.

### Flow cytometry

Differentiated cells were collected at different stages from one well of a 6-well plate. Cells were dissociated into single cells using TrypLE, washed with 1 mL of PBS and fixed with cold 4% paraformaldehyde (PFA) in PBS for 15 mins at room temperature with gentle shaking. All cell centrifugations were done at 2000 rpm for 5 mins and washes were performed using TBST (tris buffered saline with tween with added 0.5% Tween 20). Cells were permeabilized using PBST (phosphate buffered saline with 0.5% triton-X100) for 20 mins at room temperature and then blocked using 6% BSA (bovine serum albumin) in PBS containing 0.1% saponin. Blocking was done overnight at 4° C. Incubation with primary antibodies in using 3% BSA in PBS containing 0.1% saponin was done for 3 hours on a shaker, at room temperature. Cells were washed with TBST then incubated with secondary antibodies for 30 minutes at room temperature. Finally, cells were resuspended in PBS, run using BD Accuri C6 Flow Cytometer (BD Biosciences, USA) and analysed using FlowJo software. Antibodies used are listed in Supplementary Table 3.

### Immunofluorescence

Cells were fixed at different timepoints using 4% PFA in PBS for 20 min at room temperature followed by permeabilization using PBST for 15 mins. Cells were then blocked overnight at 4° C using 6% BSA in PBST with gentle shaking. For staining, these cells were incubated with primary antibodies overnight at 4° C in 3% BSA in PBST. Washes were performed with TBST. Secondary antibodies were added at room temperature for 1 hour and nuclear staining was done using Hoechst 33258 diluted 1:5000 in PBS (Life Technologies, USA). Cells were then imaged using inverted fluorescence microscope (Olympus, Tokyo, Japan). Details for antibodies used in this study are listed in Supplementary Table 3.

### Apoptosis and proliferation assays

Differentiated cells at different timepoints were dissociated using using TryplE, and were either used for apoptosis assay or fixed for proliferation assay as previously described [14]. Live dissociated cells were stained with Annexin V using Annexin V-FITC Apoptosis Detection Kit (Abcam, #ab14085) and/or 7-AAD dye (Invitrogen, Massachusetts, USA) as per manufacturer’s instructions for 5 mins at 4°C. Samples were analyzed using BD Accuri C6 Flow Cytometer and processed with FlowJo software. For proliferation assay, the cells were treated with 20 µM BrdU reagent (Invitrogen, Massachusetts, USA) for 5 hours, prior to dissociation and fixation. Dissociated single cells were then fixed with cold 70% ethanol overnight at 4°C. Cells were then rinsed once with PBS and denatured with 0.2 M HCL containing 0.5% Triton for 15 mins at room temperature, followed by neutralization with 0.1 M sodium tetraborate treatment for 20 mins. Finally, cells were washed with PBS and blocked using 6% BSA in PBS with 0.1% saponin. Cells were then incubated with Alexa Fluor 488-conjugated BrdU monoclonal antibody (B35130, Thermo Fisher, Massachusetts, USA), diluted to 1:100, at room temperature for 2 hours, washed with TBST, and BrdU^+^ cells were assessed using BD Accuri C6 Flow Cytometer and processed with FlowJo software.

### Single cell-RNA sequencing analysis

The online published GSE202497 data set for human pluripotent stem cell differentiation into pancreatic islets (https://www.ncbi.nlm.nih.gov/geo/query/acc.cgi?acc=GSE202497) was used [15] and re-analyzed as previously discussed [8] for time points D11, D14, D21, and D39. As described before, we used top 2000 highly variable genes that were obtained using Seurat V3 algorithm implement in Scanpy from a total of 25,686 cells [8]. Cells from the single-cell data were re-clustered using leiden algorithm after performing technical batch-effect correction using Harmony[cite]. Unsupervised clustering was based on the neighborhood graph computed using top 50 batch-effect adjusted principal components. Marker genes for each cluster were calculated using the Wilcoxon method implemented in rank_genes_groups function. Finally, the cell clusters were manually annotated based on expression level for marker genes and are presented in Fig. 1.

**Fig. 1:**
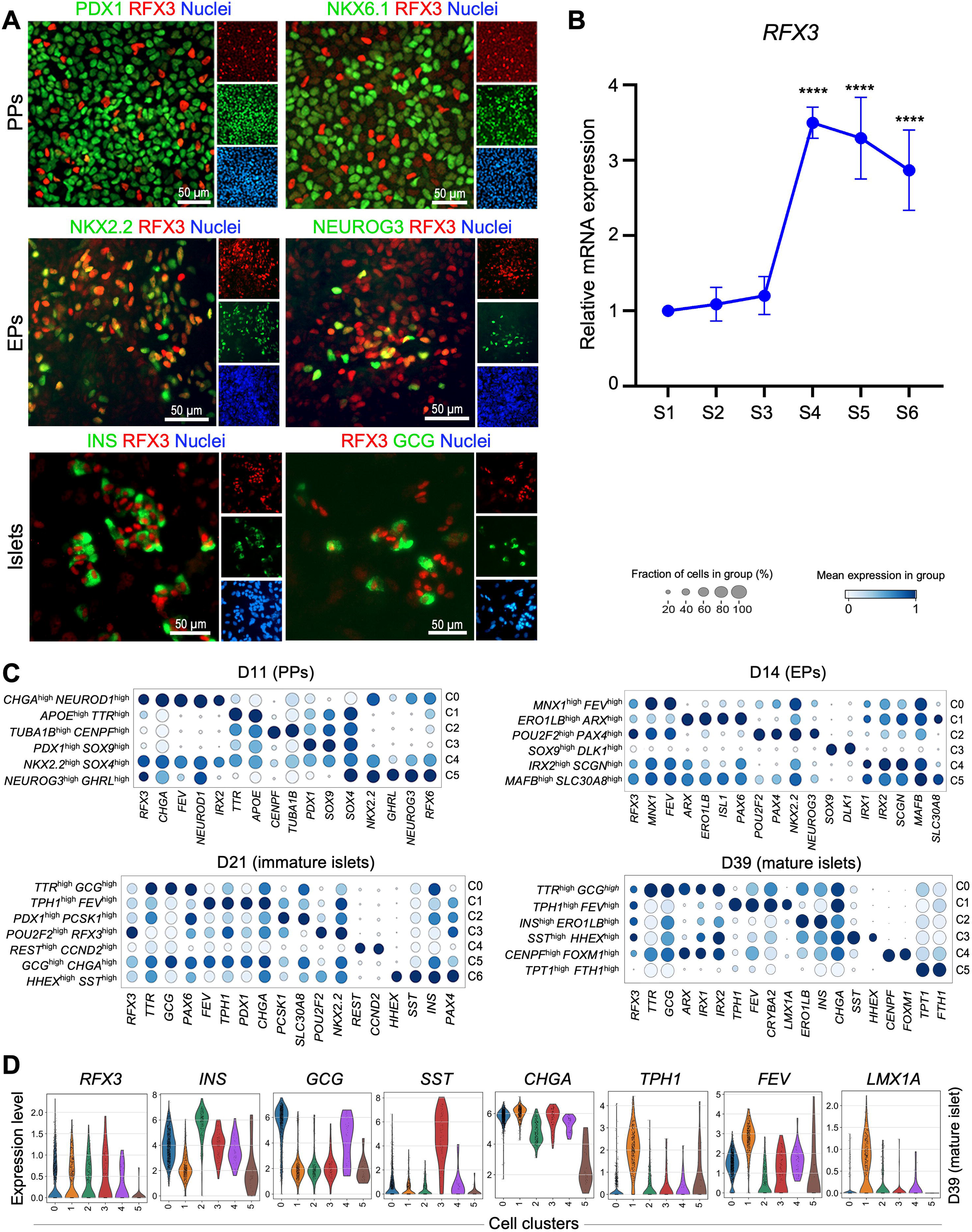
RFX3 is predominantly expressed in pancreatic endocrine lineages during differentiation of iPSCs into islets. (A) Immunofluorescence images showing stage-specific expression pattern of RFX3 in pancreatic cells. Note the absence of RFX3 expression in PDX1^+^ and NKX6.1^+^ cells at PP stage. RFX3 was co-expressed with NKX2.2 and NEUROG3 in iPSC-derived EPs and with INS and GCG in iPSC-derived islets. (B) Timeline RT-qPCR analysis for *RFX3* mRNA expression during pancreatic islet cell differentiation (n=4). (C) Dot plots demonstrating gene expression in various cell clusters determined by single-cell RNA-sequencing. Expression level in each cluster is scaled based on percentages of cells expressing *RFX3* (dot size) and mean expression (colour intensity) of the gene. Dot plots are presented for day 11 (D11), day 14 (D14), day 21 (D21), and day 39 (D39) of hESC differentiation. (D) Violin plots depicting expression pattern of key islet and enterochromaffin cell markers across various cell clusters at day 39 (mature islets). PPs, pancreatic progenitors; EPs, endocrine progenitors. The data are presented as means ±SD. \*\*\*\**p*<0.0001. Scale bar = 50 µm.

### Bulk RNA sequencing

RNA sequencing (RNA-seq) results were analyzed as previously described [8]. 1 μg of total RNA was used to purify mRNA using NEBNext Poly(A) mRNA Magnetic Isolation Kit (E7490, New England Biolabs, Massachusetts, USA). RNA-seq libraries were generated using the NEBNext Ultra Directional RNA Library Prep Kit (E7420L, New England Biolabs, Massachusetts, USA), followed by sequencing on an Illumina Hiseq 4000 system. FASTQ files were generated using Illumina BCL2Fastq Conversion Software v2.20. Pair-end FASTQ files were preprocessed with nf-core/rnaseq (version 2.7.2) pipeline, STAR (version 2.7.9a) was used for the read alignments, Salmon (version 1.5.0) for quantification of reads, TrimGalore (version 0.6.6) was applied for read trimming and GENCODE (version 38) was used for annotation of genes. Count matrix was filtered to exclude mitochondrial, ribosomal genes and low expression values were excluded using the HTSFilter (version 1.32.0) [16, 17]. Differentially expressed genes (DEGs) were identified using DESeq2 (version 1.32.0) defined by log2 fold change (FC) > 1 and <-1, with adjusted *p value* < 0.05 and pathway analysis was done using the Database for Annotation, Visualization, and Integrated Discovery (DAVID) [18].

### Glucose-stimulated insulin secretion assays

Mature islet organoids were washed with PBS, serum-starved in glucose-free Krebs buffer for 2 hours at 37° C, 5% CO_2_, and then equal numbers of islet organoids were treated with low glucose (2 mM) and high glucose in Krebs (20 mM) for 30 mins, with supernatants collected after each treatment. To measure insulin content, islet organoids were treated with 30 mM KCl in low glucose Krebs for 30 mins followed by starvation. Insulin levels in the supernatants were measured using ELISA (ELH-Insulin, RayBiotech, Georgia, USA) and normalized to total cell number in each condition.

### Western blotting

Total protein was extracted from 1-2 wells of a 6-well plate using RIPA lysis buffer containing a protease inhibitor (ThermoFisher Scientific, Massachusetts, USA). The protein concentration was measured using the Pierce BCA kit (ThermoFisher Scientific, #23225, Massachusetts, USA). Around 20 µg of total protein was loaded onto 7.5-10% SDS-PAGE gels for separation and then transferred to PVDF membranes (ThermoFisher Scientific, #88518, Massachusetts, USA). The membranes were blocked with 15% skimmed milk in TBST for at least 3 hours at room temperature or overnight at 4°C. A primary antibody was added and incubated overnight at 4°C. Following this, the membranes were washed with TBST, and a secondary antibody was applied for 1 hour at room temperature, followed by additional washes with TBST. Membranes were developed using SuperSignal West Pico Chemiluminescent substrate (ThermoFisher Scientific, #34580, Massachusetts, USA). Details of the antibodies used are provided in Supplementary Table 3.

### Statistical Analysis

At least 3 biological replicates were analysed for each experiment, and *p* values were calculated using an unpaired two-tailed Student’s *t* test on GraphPad Prism 8 software. (GraphPad Software, Boston, MA, USA, www.graphpad.com). Data are represented as mean ± SD.

## Results

### RFX3 is highly expressed in stem cell-derived pancreatic endocrine lineages

To investigate the stage-specific role of RFX3 in human pancreatic islet development, we differentiated wild type iPSCs (WT iPSCs) into pancreatic islets using an optimized protocol. We evaluated RFX3 expression at various stages of this differentiation (Fig. 1A, B, Supplementary Fig. 1). The expression of RFX3 was not detected during the definitive endoderm (DE; S1), primitive gut tube (PGT; S2), and posterior foregut (PF; S3) stages of differentiation (Supplementary Fig. 1B). However, immunofluorescence analysis showed a high expression of RFX3 in pancreatic progenitors (PPs; S4). Interestingly, RFX3 expression was notably high in PPs that did not express PDX1 and NKX6.1 (Fig. 1A). In iPSC-derived endocrine progenitors (EPs; S5), we observed co-localization of RFX3 with the endocrine markers NKX2.2 and NEUROG3. Upon differentiation to islets, all INS- and GCG-expressing cells showed nuclear expression of RFX3; however, RFX3 expression was not restricted to hormone-expressing islet cells (Fig. 1A). Furthermore, RT-qPCR analysis demonstrated that RFX3 mRNA expression increased during endocrine differentiation with the highest level being in PP, EP and islet stages (Fig. 1B).

To decipher cell type-specific role of RFX3 during pancreatic development, we re-analyzed a previously published single-cell RNA sequencing (scRNA-seq) dataset for hESC-derived pancreatic islets (GSE202497) [15]. We focused on the time points defined by the dataset as D11, D14, D21, and D39, which correspond to PPs, EPs, immature islets, and mature islets, respectively. Cell-clusters obtained after unsupervised clustering were visualized using Uniform Manifold Approximation and Projection (UMAP) were nomenclated using key mRNA expression signatures for each differentiation stage (Fig. 1C, D). At D11 (PPs), *RFX3* was strongly expressed in endocrine cell populations represented by a high proportion of *CHGA, FEV, NEUROD1, NKX2.2, NEUROG3, RFX6, GHRL, IRX2,* and *SOX4* expression within different clusters; whereas only low expression of *RFX3* was observed in multipotent progenitor cell populations represented by high expression of *PDX1* and *SOX9* (Fig. 1C). The lack of *RFX3* expression in the *PDX1*^high^/*SOX9* ^high^ cluster aligns with our immunostaining results, which also showed exclusion of RFX3 expression from PDX1^+^ and NKX6.1^+^ PPs (Fig. 1A). We observed low levels of *RFX3* expression in the *APOE*^high^/*TTR*^high^ and proliferative *TUBA1B*^high^/*CENPF*^high^ clusters (Fig. 1C).

On day 14 (EPs), *RFX3* expression was observed in most clusters that represented EP subpopulations, with the highest found in the *POU2F2*^high^/*PAX4*^high^ cluster. This cluster also showed high expression of *NEUROG3* and *NKX2.2* as well as moderate levels of *MNX1* and *FEV* (Fig. 1C). Other key endocrine markers that were highly expressed in clusters that showed moderate *RFX3* levels were *ARX*, *ISL1*, *ERO1LB*, *PAX6*, *SCGN*, *IRX1, IRX2, MAFB,* and *SLC30A8* while lowest expression of *RFX3* was observed in *SOX9*^high^/*DLK1*^high^cluster (Fig. 1C). On day 21 (immature islets), the highest expression of *RFX3* was seen in an endocrine cluster characterized by *POU2F2*^high^/*RFX3*^high^. Other endocrine clusters showed moderate levels of *RFX3* expression (Fig. 1C). Finally, on D39 (mature islets), different mature endocrine cell populations were well segregated: *INS*^high^*/ERO1LB*^high^ (beta cells), *TTR*^high^/*GCG*^high^ (alpha cells), *SST*^high^/*HHEX*^high^ (delta cells), and *TPH1*^high^/*FEV*^high^ (enterochromaffin-like cells). *RFX3* expression was prominently observed in all of these cell clusters (Fig. 1C, D). *RFX3* expression was low in the *CENPF*^high^/*FOXM1*^high^ cluster, and negligible in the *TPT1*^high^/*FTH1*^high^ cluster (Fig. 1C). These results indicate that *RFX3* is expressed during pancreatic progenitor differentiation and endocrine specification in humans, as well as in hormone-expressing mature islets. This suggest that RFX3 may be involved in regulating human pancreatic islet differentiation and the development of endocrine pancreas.

### *RFX3* deletion specifically impairs endocrine gene regulatory network in iPSC-derived pancreatic progenitors

To investigate the functional role of RFX3 in human islet development, we generated *RFX3* mutant iPSCs by targeting exon 3 to introduce indels using CRISPR/Cas9 in WT iPSCs. Resulting frameshift mutations led to introduction of a premature stop codon in exon 4. Successfully edited clones were, thus, selected following validation using sanger sequencing (Supplementary Fig. 2A). Furthermore, upon differentiation to PP stage, the generated *RFX3* KO iPSC lines showed complete absence of RFX3 expression by Western blotting and immunofluorescence (Fig. 2A, B). The selected *RFX3* KO iPSCs expressed key pluripotency markers and had normal karyotypes, comparable to those of WT iPSCs (Supplementary Fig. 2B, C). We then differentiated these *RFX3* KO iPSC lines into pancreatic islets to evaluate the effect of *RFX3* loss across stages S4 (PPs), S5 (EPs), and S6 (islets) that exhibited RFX3 expression. Our analysis revealed that PPs lacking *RFX3* exhibited no significant change in the protein and mRNA levels of key progenitor markers, such as PDX1, SOX9, and FOXA2 (Fig 2B-E). However, NKX6.1 expression was significantly reduced at protein and mRNA levels (Fig. 2B, C, E). RT-qPCR analysis showed no significant change in the mRNA levels of other progenitor markers, *ONECUT1* and *ONECUT2* (Fig. 2E).

**Fig. 2:**
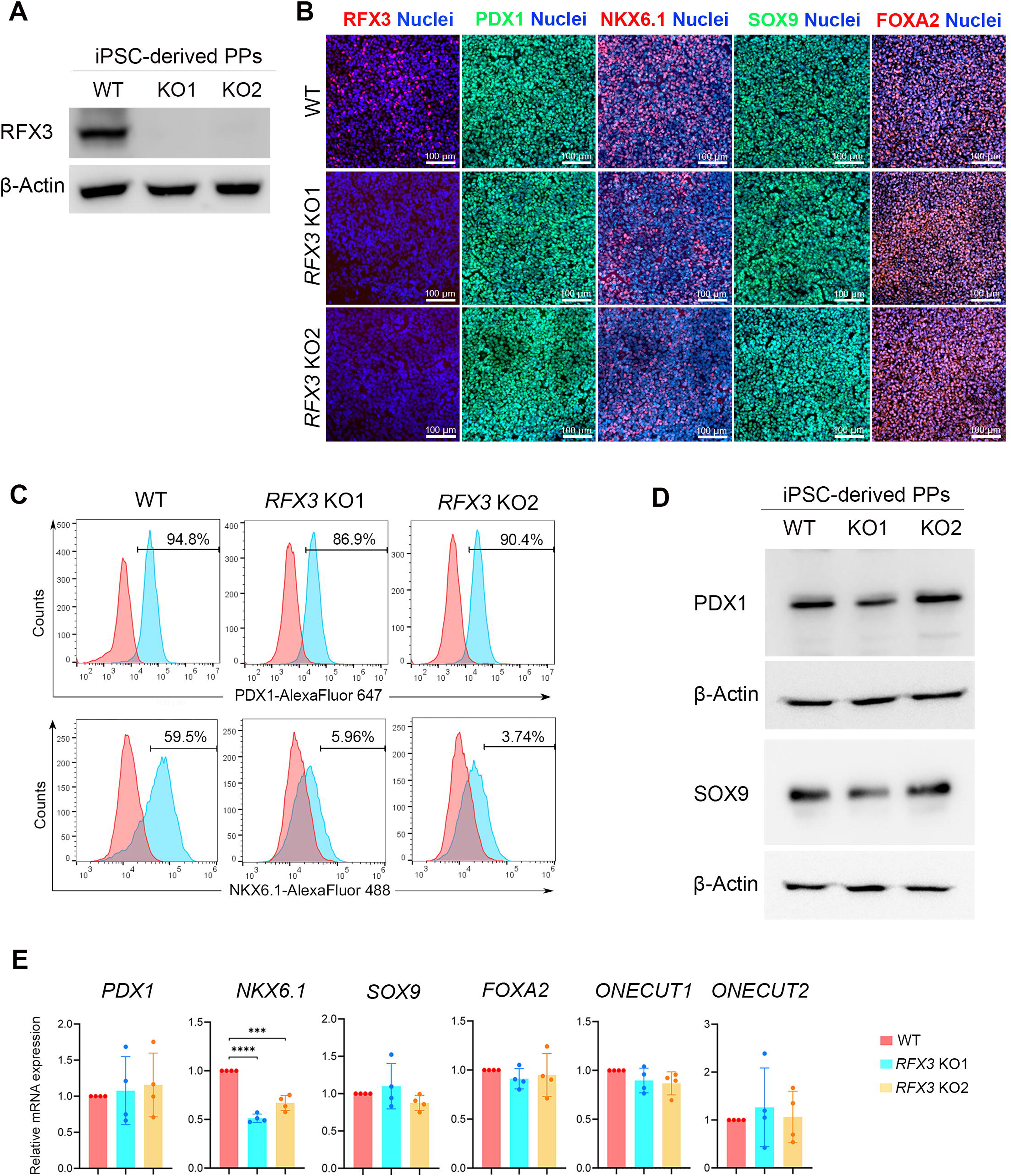
Impact of *RFX3* deletion on iPSC-derived pancreatic progenitor (PPs). (A) Western blotting analysis validating the loss of RFX3 protein expression in PPs expressing PDX1. (B) Immunofluorescence images showing expression of RFX3, PDX1, NKX6.1, SOX9, and FOXA2 in *RFX3* KO PPs and WT PPs. (C) Representative flow cytometry histograms showing the expression of PDX1 and NKX6.1 in *RFX3* KO PPs and WT PPs. (D) (E) Representative Western blotting analysis showing the protein expression levels of PDX1 and SOX9 in *RFX3* KO PPs and WT PPs. (E) RT-qPCR analysis for PP developmental markers (n=4). The data are presented as means ±SD. \*\*\**p*<0.001, \*\*\*\**p*<0.0001. Scale bar = 100 µm.

To gain further insight into the impact of *RFX3* loss on pancreatic gene network, we conducted bulk RNA-seq on *RFX3* KO PPs and WT PPs. The transcriptome analysis revealed significant dysregulation in endocrine specification processes in *RFX3* KO PPs. We identified 387 downregulated differentially expressed genes (DEGs) (log_2_ FC < -1.0, *p* value < 0.05) and 125 upregulated DEGs (log_2_ FC > 1.0, *p* value < 0.05) in *RFX3* KO PPs compared to WT PPs (Fig. 3A, Supplementary Tables 4-5, Supplementary Fig. 3A). Gene ontology analysis of the downregulated biological processes in *RFX3* KO PPs showed significant enrichment related to insulin secretion, potassium and calcium ion transport, pancreatic islet cell and endocrine pancreas development, and response to hypoxia (Fig. 3B, Supplementary Fig. 3B and 4). However, pathways related to cholesterol and triglyceride homeostasis, negative regulation of cell apoptotic process, and cellular oxidant detoxification were upregulated in *RFX3* KO PPs, indicating a shift in lineage specification in *RFX3* KO PPs (Supplementary Fig. 3B). The heatmaps for the selected top significantly downregulated DEGs governing islet development and insulin secretion suggest a disrupted endocrine program in PPs lacking *RFX3* (Fig. 3C). Furthermore, RT-qPCR validation confirmed that several key transcriptional regulators essential for islet development such as *ARX, PAX6, NKX2.2, NEUROD1, NEUROG3, RFX6, CHGA, CHGB, CRYBA2, ERO1B (ERO1LB), MAFB, PTPRN2, IRX1, IRX2, SCG3, PCSK1, PCSK2*, *INSM1, ISL1, FFAR2, FEV*, and *LMX1B* were significantly downregulated in *RFX3* KO PPs (Fig. 3D). Interestingly, comparison with recent RNA-seq results from *RFX6* KO PPs [8] revealed that only 32% of the downregulated and 26.4% of the upregulated genes in *RFX3* KO PPs were also downregulated and upregulated in *RFX6* KO PPs[8], respectively (Supplementary Fig. 5).

**Fig. 3:**
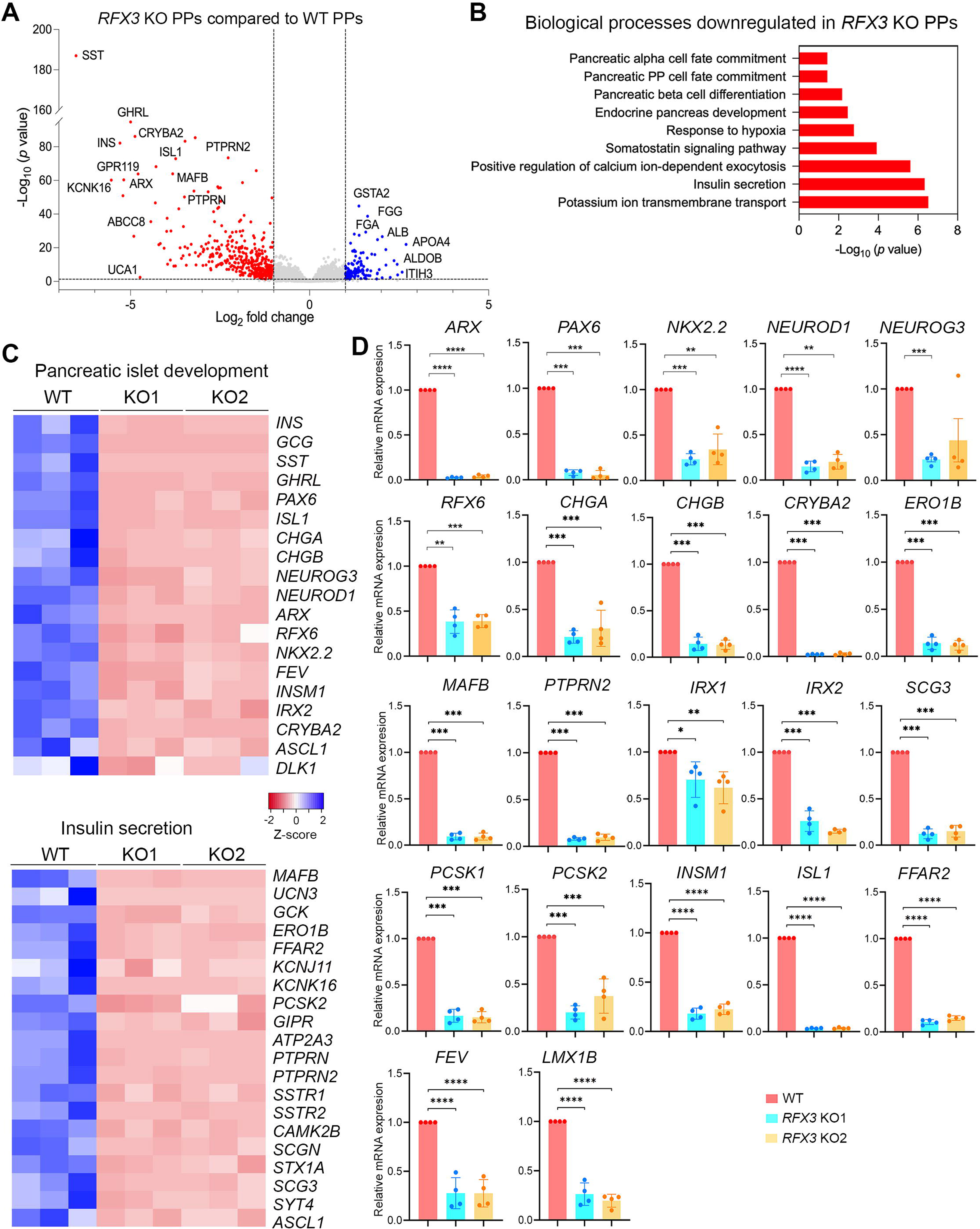
Abolished endocrine specification in iPSC-derived pancreatic progenitors lacking *RFX3*. Differentially expressed genes (DEGs) and pathways were identified by bulk RNA-sequencing of *RFX3* KO PPs and WT PPs (n=3). (A) Volcano plot showing up-(blue) and downregulated (red) DEGs. (B) Gene ontology of significantly enriched biological processes in downregulated DEGs. (C) Heatmaps depicting Z score values of significantly downregulated DEGs regulating pancreatic islet development and insulin secretion. (D) RT-qPCR analysis of key endocrine markers affected by *RFX3* deletion in iPSC-derived PPs (n=4). The data are presented as means ±SD. \**p*<0.05, \*\**p*<0.01, \*\*\**p*<0.001, \*\*\*\**p*<0.0001.

### *RFX3* deficiency reduces pancreatic endocrine genes and increases enterochromaffin cell genes in iPSC-derived endocrine progenitors

Next, we investigated the impact of *RFX3* loss on EPs. Immunofluorescence analysis of iPSC-derived EPs showed no significant differences in the protein levels of CHGA, NEUROG3, NKX2.2, and NKX6.1 compared to WT (Fig. 4A). RT-qPCR results were comparable, except for *NEUROG3*, which showed a significant increase in mRNA levels (Fig. 4B). However, several key genes associated with endocrine islet development were significantly downregulated in EPs lacking *RFX3*, including *ARX, IRX1, IRX2, PAX6, ERO1B, CRYBA2, KCNJ11, SLC30A8, IAPP, UCN3, INS, GCG, SST,* and *PPY* (Fig. 4C). Of note, scRNA-seq analysis at EP stage showed high *RFX3* expression in cell clusters expressing high levels of these endocrine genes (Fig. 1C), suggesting their direct regulation by *RFX3*. Interestingly, genes linked to EC development, such as *PAX4, FEV, CDX2, TPH1, SLC18A1,* and *LMX1A*, were significantly upregulated (Fig. 4C). The increase in the CDX2 expression was validated using Western blotting (Fig. 4D). These findings indicate that RFX3 is crucial for the proper lineage specification of endocrine islet cells.

**Fig. 4:**
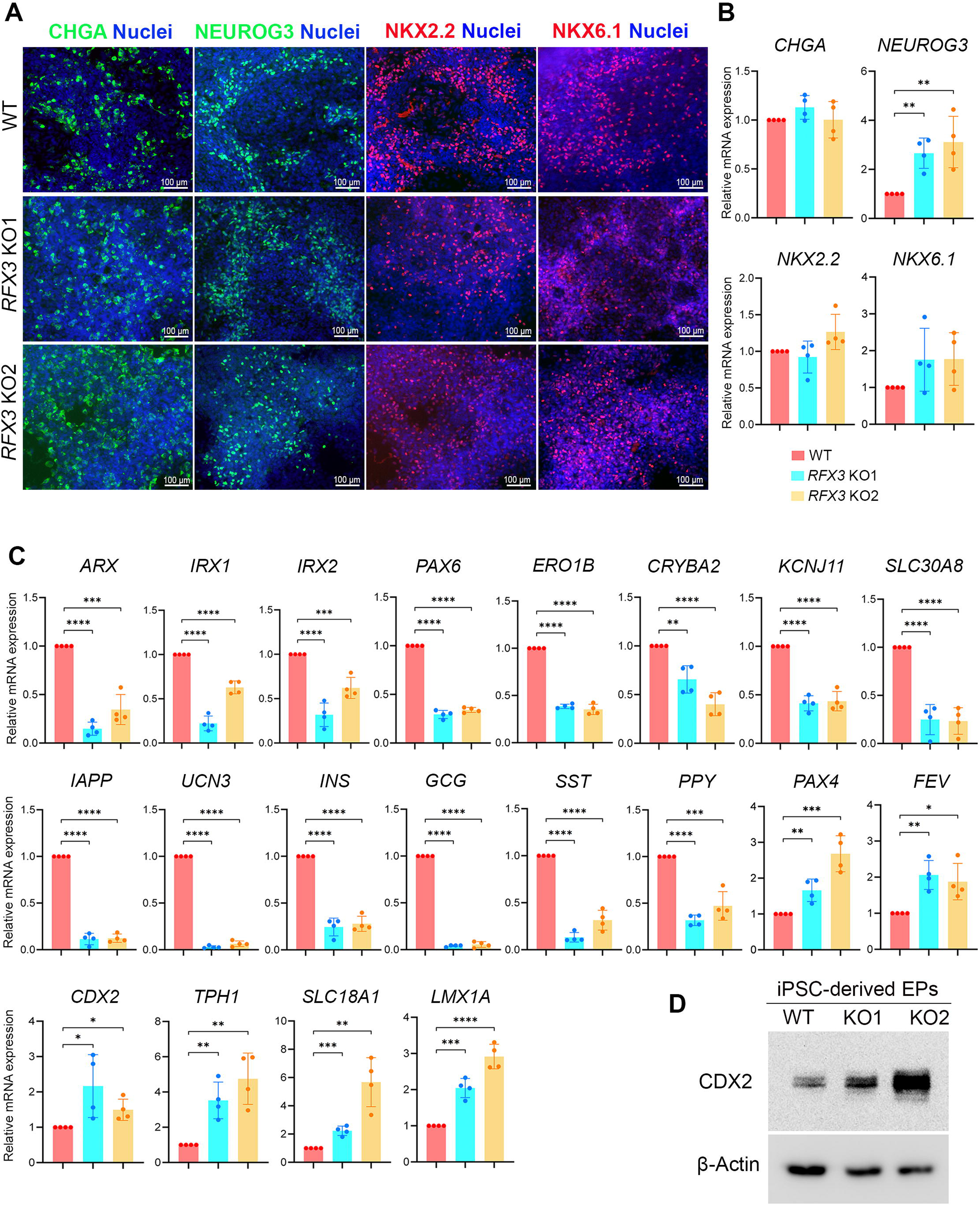
Effect of *RFX3* deletion on endocrine progenitors derived from iPSCs. Immunofluorescence images (A) and RT-qPCR (B) showing the expression of endocrine progenitor markers, including *CHGA, NKX2.2,* and *NKX6.1* in endocrine progenitors (EPs) derived from WT and *RFX3* KO iPSCs showing no significant differences. (C) RT-qPCR analysis showing reduced expression of key markers regulating endocrine pancreas differentiation, along with increased expression of enterochromaffin cell markers in EPs derived from *RFX3* KO EPs compared to WT EPs (n=4). (D) Western blotting analysis showing an increase in the expression levels of CDX2 in *RFX3* KO EPs compared to WT EPs. The data are presented as means ±SD. \**p*<0.05, \*\**p*<0.01, \*\*\**p*<0.001, \*\*\*\**p*<0.0001. Scale bar = 100 µm.

### Loss of *RFX3* disrupts the development of iPSC-derived pancreatic islets

To assess the impact of *RFX3* absence on endocrine pancreas formation and functionality, we evaluated iPSC-derived islet differentiation using different approaches. Immunofluorescence and RT-qPCR analyses revealed reduced expression of key islet hormones including INS, GCG, SST, GHRL, PPY and UCN3, while CHGA expression remained unaffected (Fig. 5A, B). Interestingly, FEV, an EC marker, was upregulated in *RFX3* KO islets (Fig. 5A). Flow cytometry quantification showed a decreased percentage of INS^+^/NKX6.1^+^ co-expressing mono-hormonal beta cells in *RFX3* KO islets, compared to WT islets (Fig. 5C).

**Fig. 5:**
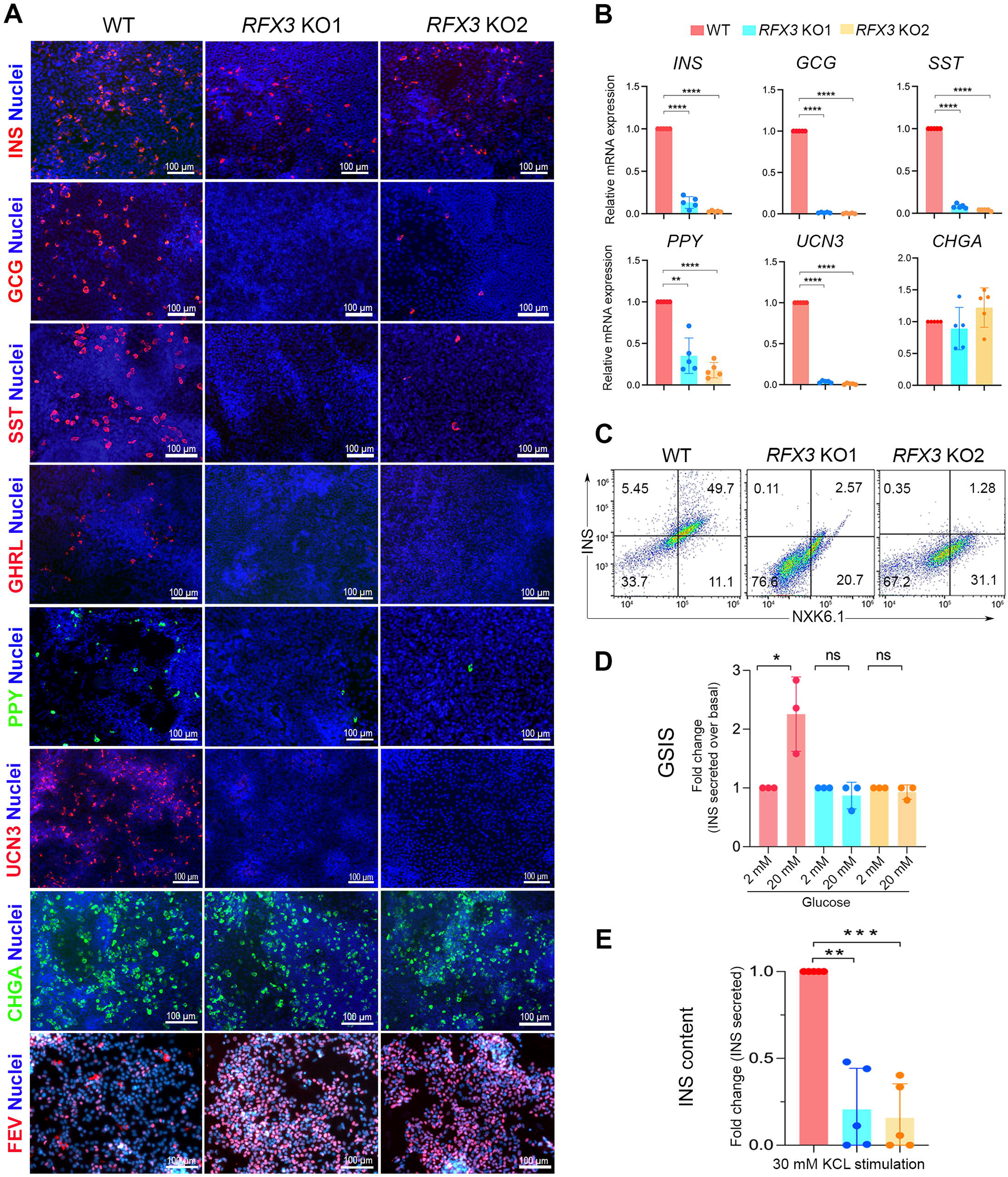
Impaired generation of hormone-secreting islet cells from *RFX3* KO iPSCs. (A) Immunofluorescence images showing reduced expression of islet hormones (INS, GCG, SST, GHRL, and PPY) and UCN3 in islet cells differentiated from *RFX3* KO iPSCs. In contrast, CHGA expression remained unchanged, while FEV expression increased. (B) RT-qPCR analysis for key islet markers in WT islets and *RFX3* KO islets (n=5). (C) Flow cytometry quantification showing decreased generation of INS^+^/NKX6.1^+^ mono-hormonal pancreatic beta cells in islets differentiated from *RFX3* KO iPSCs compared to WT controls. Glucose-stimulated insulin secretion (GSIS) assay illustrating diminished beta cell functionality (fold change in insulin secretion) (D) and reduced insulin content (E) upon treatment with high glucose (20 mM, n=3) and KCl (30 mM, n=5) respectively. The data are presented as means ±SD. \**p*<0.05, \*\**p*<0.01, \*\*\**p*<0.001, \*\*\*\**p*<0.0001. Scale bar = 100 µm.

To enhance the functional properties of iPSC-derived islet cells, we re-aggregated iPSC-derived PPs into 3D organoids and differentiated them into islet cells. We then performed glucose-stimulated insulin secretion (GSIS) assays on these iPSC-derived islet organoids to explore the impact of *RFX3* deficiency. Upon treatment with high glucose concentrations (20 mM), *RFX3* KO islet organoids showed an abolished insulin secretion response compared to WT organoids (Fig. 5D). This indicates that *RFX3* depletion impairs the functionality of pancreatic beta cells by diminishing their insulin secretion capacity. Furthermore, stimulation with the depolarizing agent KCl failed to induce an adequate insulin secretion response from *RFX*3 KO islet organoids, indicating a significantly reduced insulin content (Fig. 5E).

To gain a deeper understanding of the role of RFX3 in human islets, we performed RNA-seq on *RFX3* KO islets and WT islets. The transcriptomic data revealed hampered islet development due to the dysregulation of key endocrine genes (Fig. 6A-C; Supplementary Fig. 3). We identified 557 DEGs that were significantly downregulated (log_2_ FC < -1.0, *p* value < 0.05), and 383 DEGs that were upregulated (log_2_ FC >1.0, *p* value < 0.05) in *RFX3* KO islets compared to WT islets (Supplementary Tables 6-7). Islet hormones including *INS, GCG, SST, PPY,* and *GHRL* were downregulated, consistent with the results from immunostaining and qPCR (Fig. 6C, D). Furthermore, genes directly targeted by RFX3, such as *GCK*[5], were also downregulated, along with several other key markers of pancreatic endocrine differentiation (Fig. 6A, C-D). Gene ontology analysis of downregulated DEGs revealed significant enrichment of pathways related to insulin secretion, potassium and calcium ion transport, response to hypoxia, among others associated with islet development and function (Fig. 6B, C; Supplementary Fig. 4). However, amongst the significantly upregulated processes, liver and fat cell development, proteolysis, triglyceride biosynthesis, lipid and glycerol metabolic processes, and urea metabolism were enriched (Supplementary Fig. 3B). RT-qPCR validation confirmed a significant downregulation in the expression of pancreatic endocrine genes, including *GCK, PAX6, ARX, ISL1, IRX1, IRX2, SIX3, ERO1B, FFAR1, MAFB, IAPP, KCNJ11,* and *SLC30A8* (Fig. 6D). RT-qPCR also confirmed the upregulations of *APOC3* and *TXNIP,* a critical gene for pancreatic islet survival [19] (Fig. 6D). Interestingly, the EC markers *FEV, TPH1, SLC18A1, LMX1A,* and *CDX2* were significantly upregulated in *RFX3* KO islets compared to WT controls (Fig. 6D). Our findings indicate that RFX3 is essential for the formation of pancreatic islets and for suppressing the development of ECs, which is linked to beta cell immaturity [15].

**Fig. 6:**
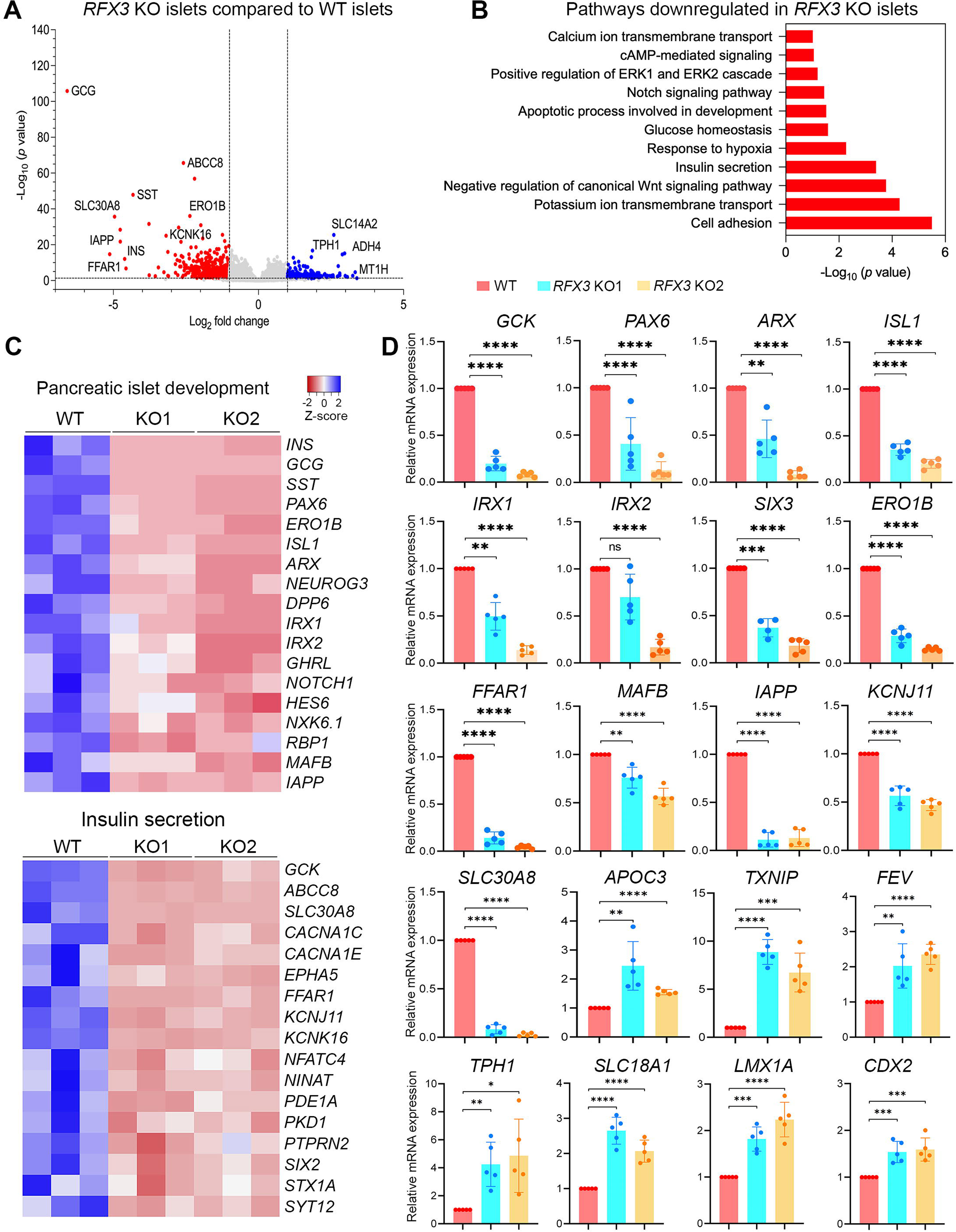
Transcriptome changes in pancreatic islets lacking the *RFX3* gene. (A) Volcano plots depicting differentially expressed genes (DEGs) identified by transcriptome analysis of islet cells derived from *RFX3* KO iPSCs and WT iPSCs (n=3). Significantly upregulated DEGs are shown in blue while downregulated DEGs are in red. (B) Gene ontology of downregulated biological processes in *RFX3* KO islets compared to WT islets. (C) Heatmaps for key DEGs involved in pancreatic islet development and insulin secretion depicting their Z score values. (D) RT-qPCR analysis showing downregulation of key islet markers and upregulation of genes associated with enterochromaffin cells in iPSC-derived islets lacking *RFX3* (n=5). The data are presented as means ±SD. \**p*<0.05, \*\**p*<0.01, \*\*\**p*<0.001, \*\*\*\**p*<0.0001.

### *RFX3* deficiency reduces the viability of iPSC-derived islet organoids

During differentiation of islet organoids, we observed dramatic morphological differences between *RFX3* KO and WT organoids (Fig. 7A). In *RFX3* KO organoids, the structure deteriorated into smaller clumps of dissociating cells during stage 5 of differentiation, and this abnormal morphology persisted into stage 6, contrasting with the stable morphology observed in WT organoids (Fig. 7A). To determine whether the reduced size of islet organoids is due to increased apoptosis or decreased cell proliferation, we assessed both processes. Flow cytometry analysis revealed a significant increase in Annexin V^+^ (apoptotic) cells in pancreatic cells derived from *RFX3* KO iPSCs at both PP and EP stages (Fig. 7B). Furthermore, quantifying BrdU incorporation showed no significant difference in proliferation rates between WT and KO cells, indicating no impairment in proliferation capacity (Fig. 7C). To explore the mechanism of cell death, we analyzed TXNIP protein levels, which are essential for islet survival[19] and were found to be increased in our RNA-seq and qPCR analyses. Western blotting confirmed a significant increase in TXNIP in EPs lacking RFX3 compared to WT controls (Fig. 7D). These results suggest that RFX3 loss impairs islet organoid viability during development by increasing TNXIP levels.

**Fig. 7:**
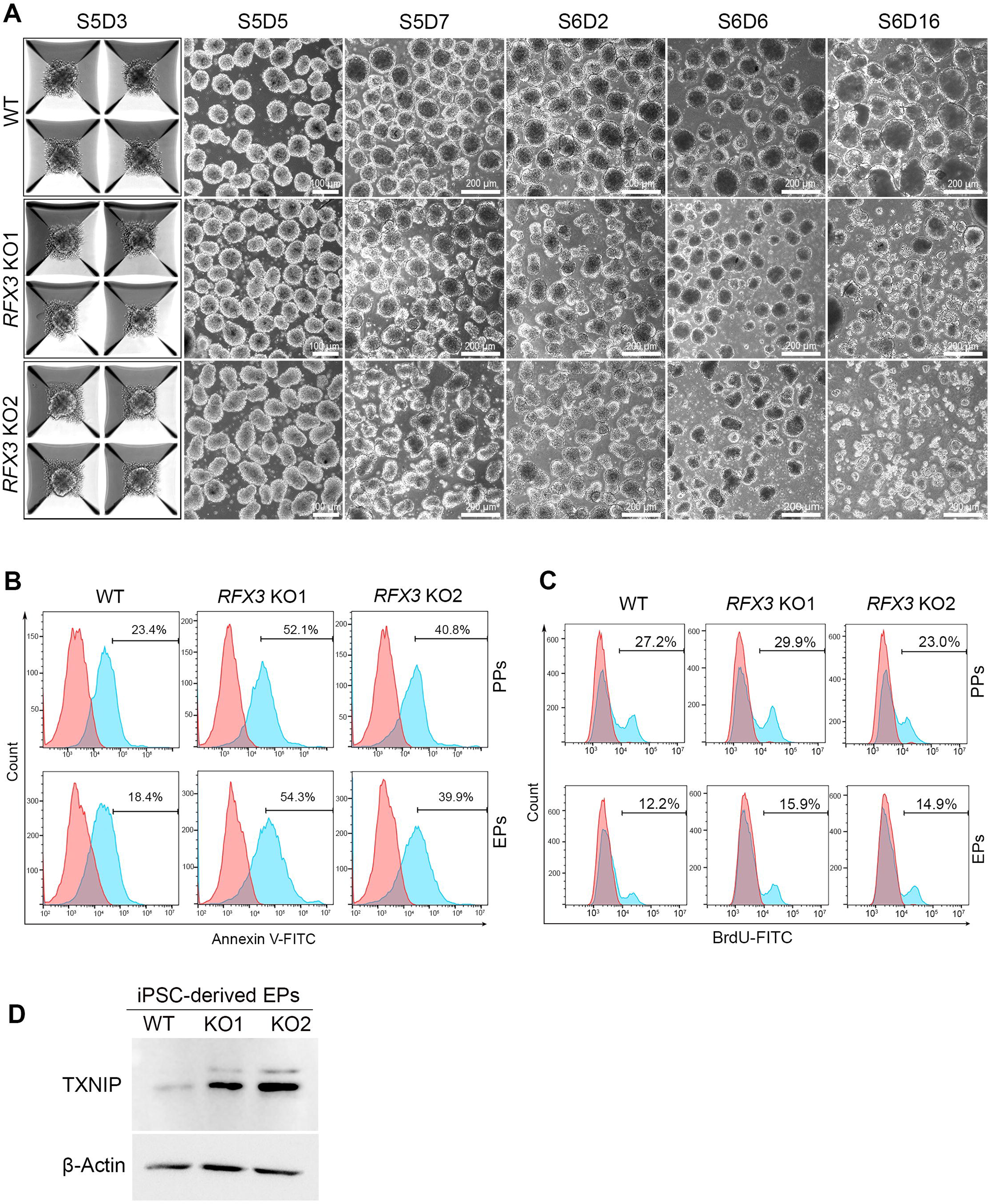
*RFX3* loss leads to increased apoptosis and disruption of iPSC-derived islet organoids. (A) Phase contrast images showing morphological changes of islet organoids during stages 5 and 6 (EPs and islets) derived from *RFX3* KO iPSCs and WT iPSCs. Note the reduced size of organoids derived from *RFX3* KO iPSCs compared to WT controls. (B) Flow cytometry quantification of apoptotic (Annexin V^+^) cells in stages 4 (PPs) and 5 (EPs) indicates increased apoptosis in *RFX3*-deficient cells. (C) Flow cytometry analysis of BrdU incorporation showing no significant changes in cell proliferation (BrdU^+^ cells) in PPs and EPs derived from *RFX3* KO iPSC lines compared to those from WT iPSCs. (D) Western blotting analysis showing an increase in the expression levels of TXNIP in *RFX3* KO EPs compared to WT EPs. Scale bars = 100, 200 µm.

## DISCUSSION

Previous rodent studies have demonstrated that RFX3 is important for the development of pancreatic islet cells [4, 5]. Recent scRNA-seq studies on hPSC-derived pancreatic islets further demonstrated that *RFX3* is expressed in multiple pancreatic endocrine cell populations at different stages of differentiation, particularly during later stages of islet development [15, 20, 21]. However, the role of RFX3 in human islet development has not yet been explored. To address this gap, we used hPSC-derived islet organoids to model *RFX3* loss-of-function and assess its impact on islet development. Our findings reveal that RFX3 expression began in PPs and was predominantly present in endocrine cell populations throughout differentiation. Loss of RFX3 significantly reduced the expression of genes critical for endocrine islet development, led to the formation of small islet organoids due to increased cell death, and impaired the development of all islet cells. Furthermore, RFX3 deficiency resulted in increased expression of enterochromaffin cell-specific genes.

Our findings are partially consistent with previous studies in rodents, which showed that *Rfx3^-/-^* mice and those with pancreas-specific *Rfx3* deletion fail to develop Ins-, Gcg- and Sst-secreting cells and show decreased Gck expression of [4, 5]. However, our results diverge from these rodent models in several key aspects. Unlike rodents, our human iPSC-derived islets did not develop PPY^+^ endocrine cells in the absence of RFX3. We also noticed significant cell death during endocrine specification in *RFX3*-deficient islet organoids, resulted in smaller islet organoids compared to WT controls. This contrasts with rodent studies that reported no reduction in the survival or proliferation in *Rfx3*-deficient cells during pancreatic development [5]. Furthermore, our human iPSC model revealed decreased expression of key endocrine transcriptional regulators, including *ARX, PAX6, NKX2.2, NEUROD1, IRX1, IRX2, CRYBA2, ERO1B, MAFB, INSM1, ISL1* and others at last three differentiation stages. Single-cell analysis further demonstrated that *RFX3* was associated with endocrine cell clusters throughout development, showing the highest expression in specific endocrine subpopulations at different stages such as *CHGA*^high^/*NEUROD1*^high^, *POU2F2*^high^/*PAX4*^high^ and *POU2F2*^high^/*RFX3*^high^ at PPs, EPs, and islets, respectively. These clusters also included key pancreatic endocrine regulators such as *CHGA, FEV, NEUROD1, NKX2.2, NEUROG3, RFX6, GHRL, IRX2, PAX6,* and *SOX4,* which support our results that RFX3 plays a role in pancreatic endocrine specification. In mature islets, *RFX3* was prominently expressed in clusters representing alpha, beta, delta, and ECs, underscoring its contribution to the complex regulatory networks governing pancreatic endocrine cell function and identity.

In this study, we observed that although the loss of RFX3 caused a significant reduction in pancreatic islet hormones, the expression of the pan-endocrine marker CHGA remained unchanged at both the EP and islet stages. The unchanged CHGA expression may be due to an increase in ECs, which also express CHGA[13]. A recent study identified three types of CHGA^+^ endocrine cells in EPs and islets derived from hPSCs: beta cells, alpha cells, and ECs[13]. Previous reports have noted the presence of ECs in fetal and hPSC-derived islets [13, 15, 20, 22], but their numbers decline with extended islet culture and are absent in the adult pancreas[15, 20]. Our results, which showed high expression levels of EC markers in EP and islet cells lacking *RFX3*, support the idea that the unchanged CHGA expression is due to a higher number of ECs, despite the reduced numbers of alpha, beta, and delta cells. We also noticed increased expression of *NEUROG3* and *PAX4*, along with unchanged levels of NKX6.1 and NKX2.2 in *RFX3* KO EPs, suggesting these markers are associated with EC development. NKX6.1, which binds LMX1A (a transcription factor for serotonin (5HT) synthesis genes) [15, 23], is reduced in pancreatic endocrine cells lacking the *CDX2* gene [15], a key regulator of EC genes like TPH1[24] and SLC18A1[15]. Zhu et al. suggested that hPSC-ECs represent a transient, beta cell-related population that diminishes with maturation, as indicated by the reduced percentage of NKX6.1^+^/SLC18A1^+^ hPSC-ECs during islet maturation [15]. In human fetal beta cells [25] and hPSC-islet cells, the 5HT-producing pre-beta cell population exhibits high expression of CHGA, SLC18A1, LMX1A, FEV, PAX4, NEUROG3, and CDX2 ^3, 15, 16, 20^. Interestingly, our results showed a significant upregulation of all these EC markers in *RFX3*-deficient EPs and islets. These findings suggest that RFX3 is essential for human islet development and maturation by promoting pancreatic endocrine specification and repressing EC differentiation.

Our study found that 32% of downregulated genes in *RFX3* KO PPs were also downregulated in *RFX6* KO PPs [8], primarily related to endocrine specification. Furthermore, 67.95% were uniquely downregulated in *RFX3* KO PPs, indicating that while RFX3 and RFX6 share some phenotypes, they also have distinct downstream mechanisms. For instance, *RFX3* deficiency increased EC markers, whereas *RFX6* loss reduced them [8]. Both *RFX3* and *RFX6* loss led to apoptosis and smaller islet organoids, but through different mechanisms. *RFX6* loss caused apoptosis due to reduced catalase (CAT) expression [8], while CAT levels remain unaffected in *RFX3*-deficient pancreatic cells. Instead, *RFX3* loss increased TXNIP expression in EPs and islets. Given that TXNIP is a pro-apoptotic protein linked to diabetes and oxidative stress-induced beta cell death [26–28], these findings suggest that *RFX3* loss impairs islet survival by elevating TXNIP levels and disrupting endocrine specification.

While *RFX6* mutations are linked to neonatal diabetes and T2D [29–33], the role of RFX3 in diabetes is less understood. Recent report identified RFX3 as part of an islet-specific enhancer complex, interacting with key genes such as GLIS3[34]. The deletion of this enhancer hub reduces RFX3 expression, underscoring its importance in islet function and its potential contribution to T2D susceptibility[34]. This highlights RFX3 importance for islet function and its possible impact on diabetes risk, but further studies are needed to understand its role in diabetes.

’In conclusion, our study highlights the essential role of RFX3 in human pancreatic islet development. We found that RFX3 is highly expressed in endocrine cells throughout islet development. Its absence impaired the formation of all pancreatic hormone-secreting cells, primarily due to decreased expression of key endocrine genes and increased cell apoptosis. This cell death was associated with elevated TXNIP levels in the absence of RFX3. Notably, RFX3 loss led to increased EC cell formation, underscoring its critical role in beta cell maturation. These results contrast with rodent studies, which did not report similar changes in apoptosis or beta cell and EC specification. Such differences may arise from technological advances or physiological differences between human and rodent islets. Our findings establish RFX3 as a crucial transcriptional regulator necessary for the proper formation and maturation of human islet cells.

## Supporting information

Suppmenetary Figures 1-5

## Contributions statement

BM, NA, AKE performed the experiments and analyzed the data. SI and SH analyzed the sequencing data. BM and EMA wrote the manuscript. EMA conceived and designed the study, supervised the project, analyzed and interpreted the data. All authors critically reviewed the article and approved the final version of the manuscript.

## Authors’ relationships and activities

S.H. is a co-founder and shareholder of Sequantrix GmbH and has research funding from by Novo Nordisk and Askbio. The authors declare that there are no relationships or activities that might bias, or be perceived to bias, their work.

