## Supplementary material for "RFX3 is essential for the development and maturation of human pancreatic islets derived from pluripotent stem cells": Suppmenetary Figures 1-5

### Supplementary Figures

#### Supplementary Figure 1

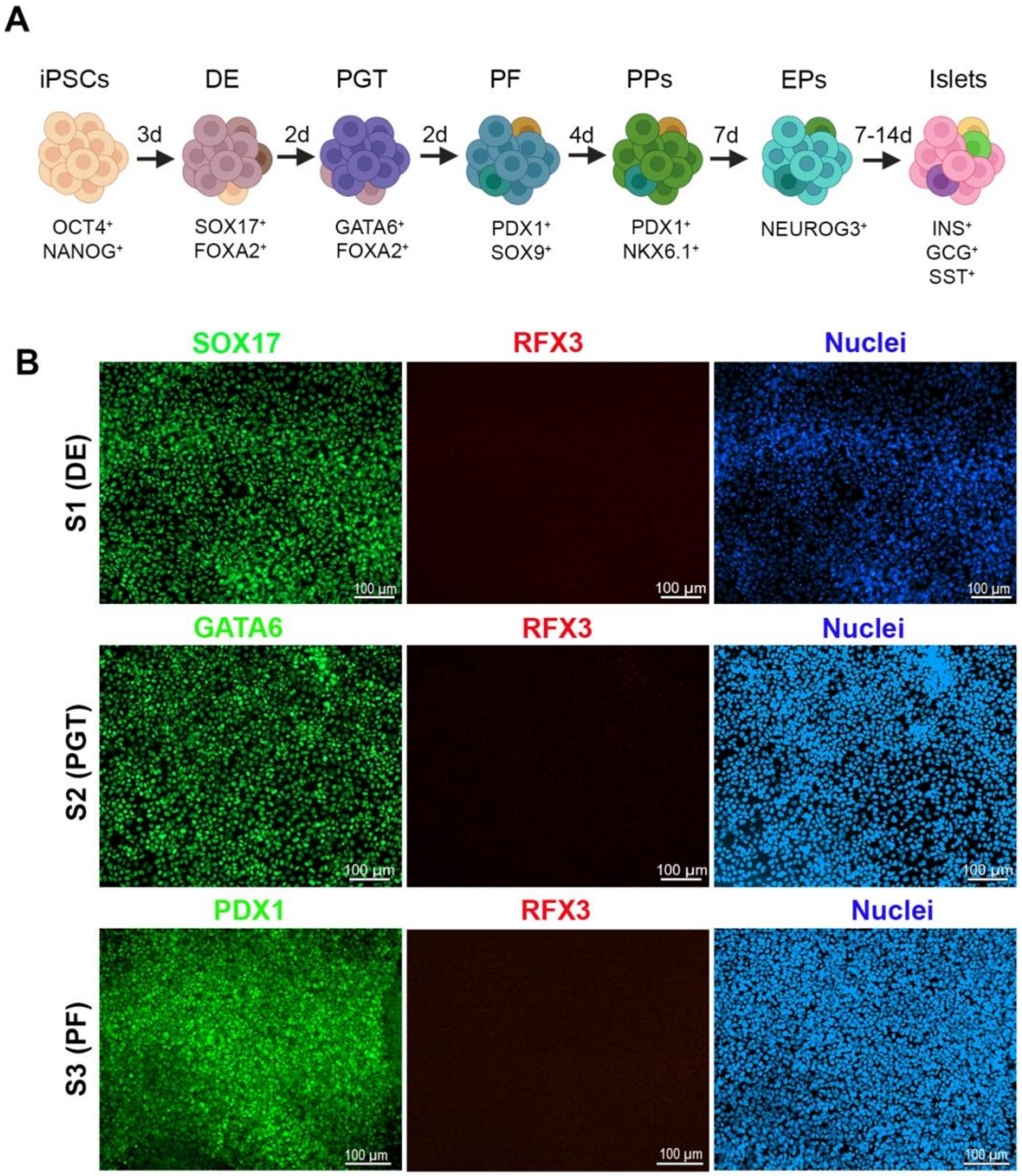


**Figure S1**: RFX3 expression at different stages during iPSC differentiation into pancreatic islets. (A) Schematic representation of the *in vitro* pancreatic islet differentiation protocol. (B) Immunofluorescence images for co-expression of RFX3 with definitive endoderm (SOX17), primitive gut tube (GATA6) and posterior foregut (PDX1) markers during early stages of iPSC differentiation to islets. Note the absence of RFX3 (red) expression in early stages of islet differentiation. Scale bar = 100 µm.

**Supplementary Figure 2**

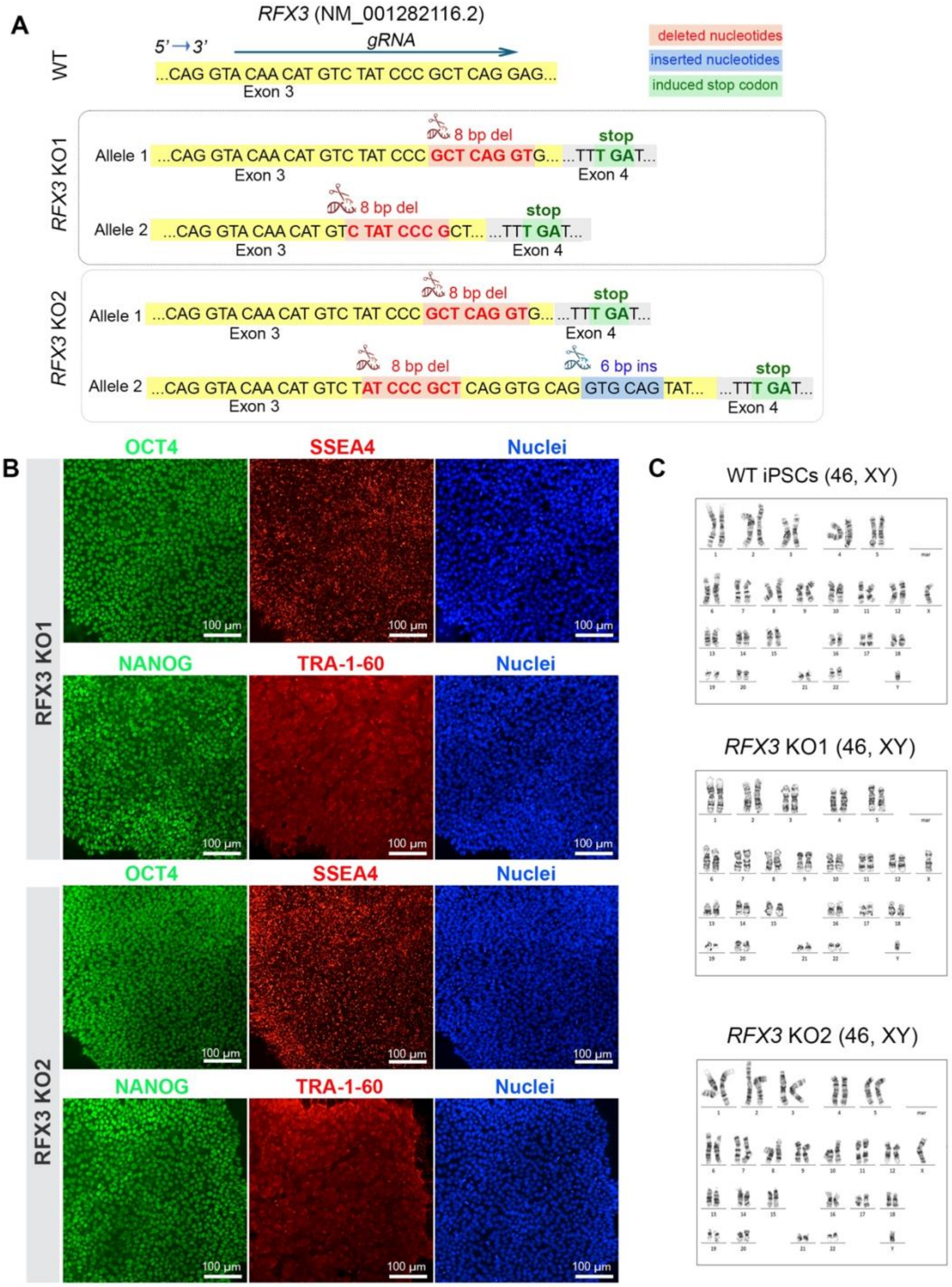


**Figure S2**: Validation and characterization of *RFX3* KO iPSC lines. (A) Sanger sequencing results demonstrating the edits and deletions in the *RFX3* gene, which resulted in the formation of stop codons. (B) Immunofluorescence images for expression of pluripotency markers for *RFX3* KO iPSCs showing their high levels in undifferentiated cells, and (C) karyotype analysis of *RFX3* KO clones showing normal number of chromosomes, similar to WT. Scale bar = 100 µm.

**Supplementary Figure 3**

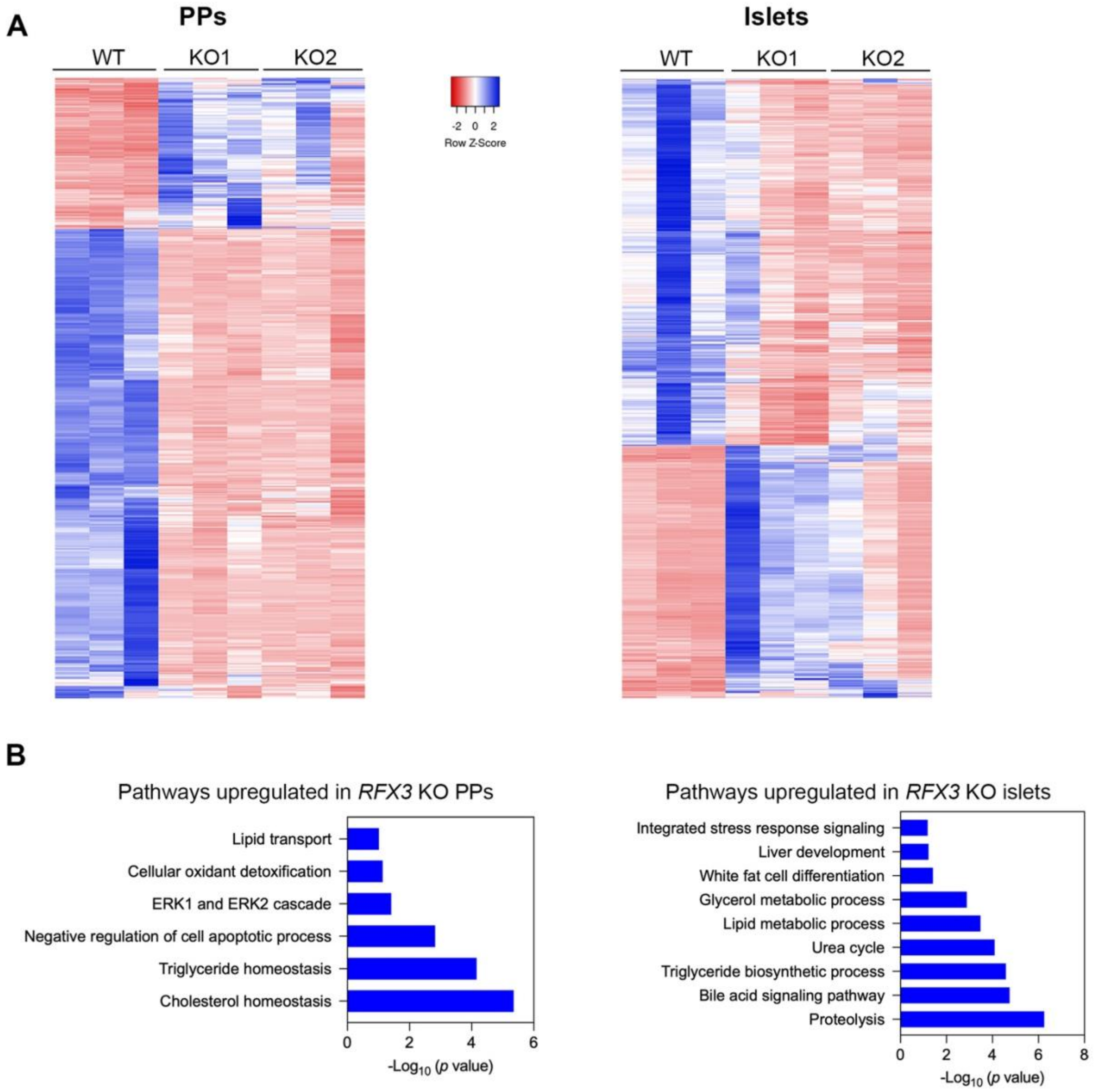


**Figure S3**: Transcriptome profiling alterations associated with *RFX3* loss in pancreatic progenitors (PPs) and islets. (A) A general clustering heatmap of differentially expressed genes (DEGs) in PPs and islets derived from WT and *RFX3* KO iPSCs. (B) Selected gene ontology pathways associated with upregulated DEGs in PPs and islets.

### Supplementary Figure 4

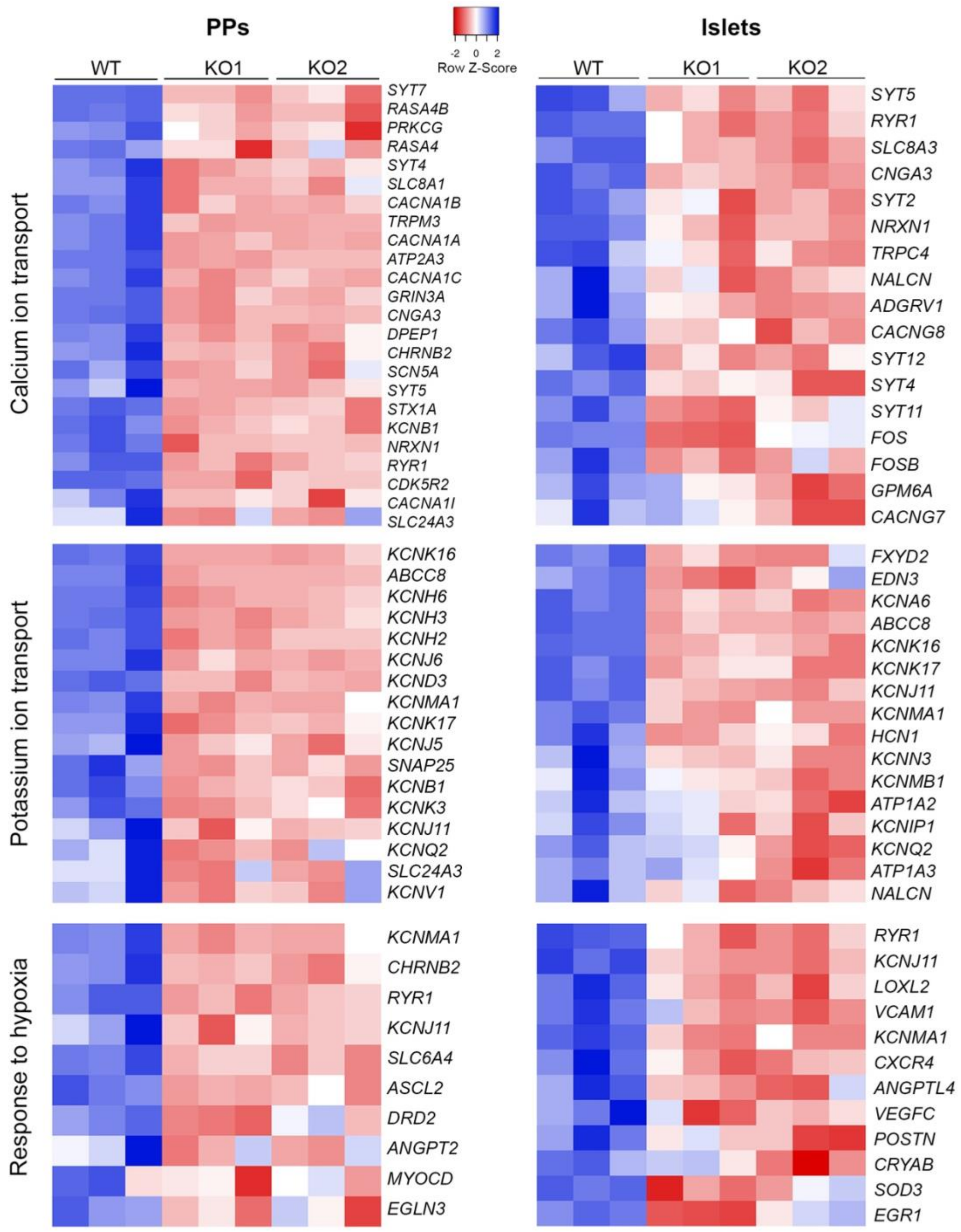


**Figure S4**: Heatmaps of downregulated DEGs highlighted enrichment in pathways related to calcium and potassium ion transport, as well as response to hypoxia, in *RFX3* KO PPs and islets, comparted to WT controls.

**Supplementary Figure 5**

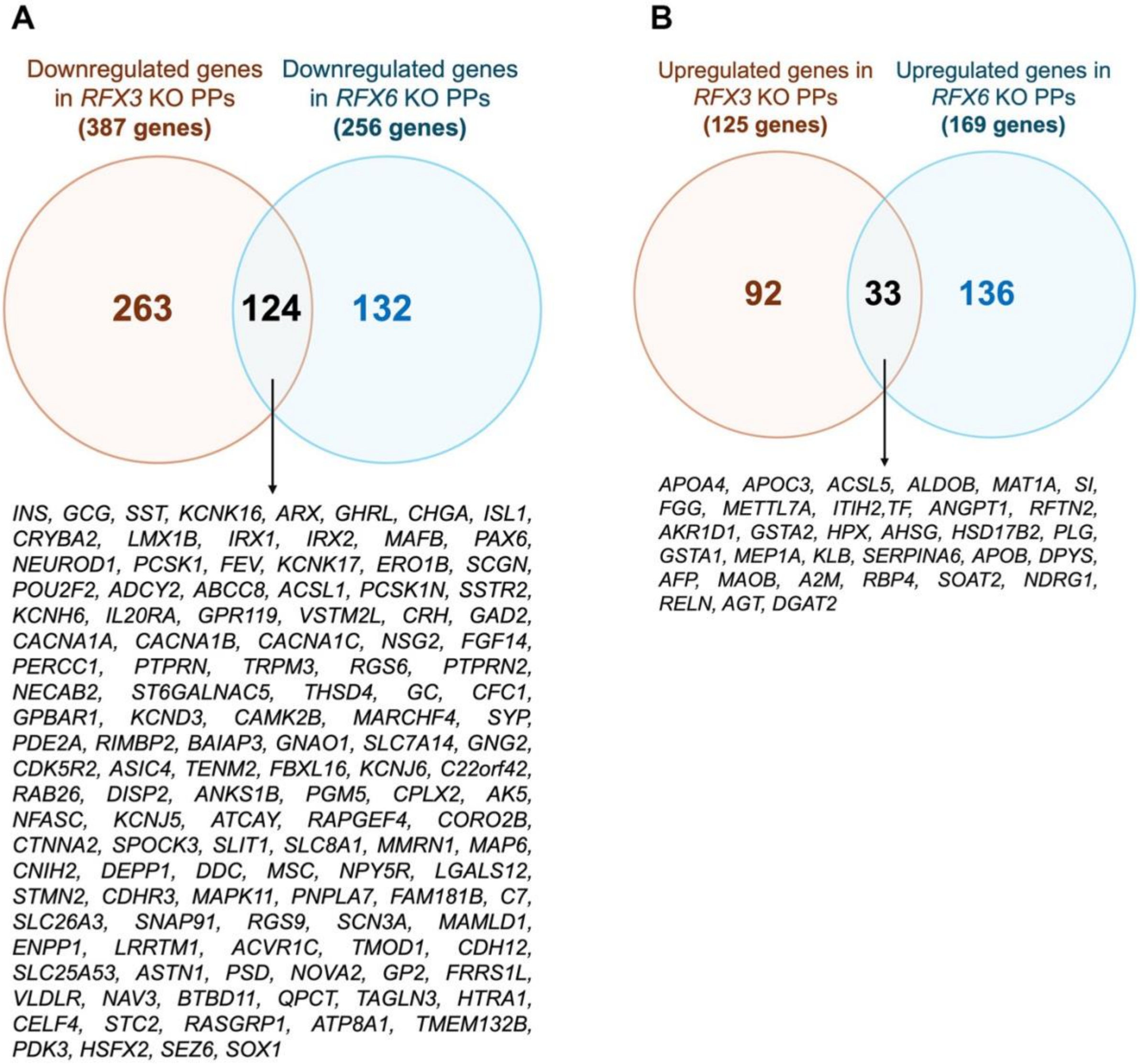


**Figure S5**: Common differentially expressed genes (DEGs) in pancreatic progenitors lacking *RFX3* and *RFX6*. (A) 124 DEGs were commonly downregulated in both *RFX3* KO and *RFX6* KO pancreatic progenitors (PPs) compared to WT PPs, representing 32% of the downregulated genes in *RFX3* KO PPs, while 263 genes (67.95%) were specically downregulated in *RFX3* KO PPs, but not in *RFX6* KO PPs. (B) A total of 33 DEGs were commonly upregulated in both *RFX3* KO and *RFX6* KO PPs compared to WT PPs, representing 26.4% of the upregulated DEGs in *RFX3* KO PPs. 92 out of 125 genes (73.6%) were specically upregulated in *RFX3* KO PPs, but not in *RFX6* KO PPs compared to WT PPs.
