## Supplementary Tables 1-7 for "RFX3 is essential for the development and maturation of human pancreatic islets derived from pluripotent stem cells"

**Table S1.** Media formulation and details of cytokine added for *in vitro* differentiation protocol.

| Differentiation stage | Media supplement | Cytokines | Total number of days |
|---|---|---|---|
| Stage 1 (DE) | **MCDB 131**:<br>1% Pen/Strep<br>1% L-Glutamine<br>0.5% Fatty acid free BSA<br>1.5 g/L $NaHCO_3$<br>10 mM D-Glucose | **Day 1**:<br>1 µM Y-27632<br>2 µM CHIR99021<br><br>**Days 1-3**:<br>100 mM Activin A<br>0.25 mM Vitamin C | 3 |
| Stage 2 (PGT) | **MCDB 131**:<br>1% Pen/Strep<br>1% L-Glutamine<br>0.5% Fatty acid free BSA<br>1.5 g/L $NaHCO_3$<br>10 mM D-Glucose | 50 ng/mL FGF-10<br>0.75 µM Dorsomorphin<br>3 ng/mL WNT-3a<br>0.25 mM Vitamin C | 2 |
| Stage 3 (PF) | **DMEM**:<br>1% Pen/Strep<br>4.5 g/L Glucose<br>110 mg/L Sodium pyruvate | 50 ng/mL FGF-10<br>200 nM LDN193189<br>0.25 µM SANT-1<br>2 µM Retinoic acid<br>0.25 mM Vitamin C<br>1% B27 w/o Vitamin A | 2 |
| Stage 4 (PPs) | **DMEM**:<br>1% Pen/Strep<br>4.5 g/L Glucose<br>110 mg/L Sodium pyruvate | 100 ng/mL EGF<br>10 mM Nicotinamide<br>200 nM LDN193189<br>0.25 mM Vitamin C<br>1% B27 w/o Vitamin A | 4-6 |
| Stage 5 (EPs) | **MCDB 131:**<br>1% Pen/Strep<br>1% L-Glutamine<br>2% Fatty acid free BSA<br>1.7 g/L NaHCO3<br>3.6 g/L Glucose<br>1:200 ITS-X100<br>10 mg/L Heparin | **Days 1-4**:<br>0.25 µM SANT-1<br><br>**Days 1-7**:<br>20 ng/mL Betacellulin<br>1 µM G-Secretase inhibitor<br>1 µM T3<br>10 µM ALK5 inhibitor II<br>0.25 mM Vitamin C<br>0.1 µM Retinoic acid | 7 |
| Stage 6 (Islet) | **MCDB 131:**<br>1% Pen/Strep<br>1% L-Glutamine<br>2% Fatty acid free BSA<br>1.23 g/L NaHCO3<br>0.45 g/L Glucose<br>1:200 ITS-X100 | 0.25 mM Vitamin C | 7-14 |

**Table S2.** List of primers used in the study.

| Gene | Forward primer | Reverse primer |
|---|---|---|
| *APOC3* | CTTCATGCAGGGTTACATGAAG | TTTCAGGGAACTGAAGCCATC |
| *ARX* | CTGCTGAAACGCAAACAGAGGC | CTCGGTCAAGTCCAGCCTCATG |
| *CDX2* | CTGGAGCTGGAGAAGGAGTTTC | ATTTTAACCTGCCTCTCAGAGAGC |
| *CHGA* | GAAGAAGGCCCCACTGTAGT | TTCCCAGCTCCATCCACAG |
| *CHGB* | CACGCCATTCTGAGAAGAGC | TCTCCTGGCTCTTCAAGGTG |
| *CRYBA2* | GATGTGGGTTCCCTCAAAGT | GCTCACCGTAAGTACAGAACTC |
| *ERO1B* | TGAACCCAGAGCGTTACACT | GGCGCCAGAGGATTTAAAGG |
| *FEV* | GCCTCTCCAAACTCAACCTC | CAAGCTGGGACTGGGGTAG |
| *FFAR1* | TGTACCCCAATCTAGGAGGC | GACCCCTTCCCAAGTAACCG |
| *FFAR2* | CGGCCTCTGTATGGAGTGAT | CCTGCTCAGTCGTGTTCAAG |
| *FOXA2* | GGGAGCGGTGAAGATGGA | TCATGTTGCTCACGGAGGAGTA |
| *GATA6* | AAGCGCGTGCCTTCATCA | TCATAGCAAGTGGTCTGGGC |
| *GAPDH* | ACGACCACTTTGTCAAGCTCATTTC | GCAGTGAGGGTCTCTCTCTTCCTCT |
| *GCG* | CTCTTCACCTGCTCTGTTCTAC | TGGATTTCTCCTCTGTGTCTTG |
| *GCK* | GCATCTTCCAGCTCTTCGAC | GGGCTACATTTGAAGGCAGA |
| *IAPP* | TTGAGAAGCAATGGGCATCC | GGGTGTAGCTTTCAGATGGTTC |
| *INS* | AAGAGGCCATCAAGCAGATCA | CAGGAGGCGCATCCACA |
| *INSM1* | TTTGTCTCGTGGTTGGAAGC | CCAAAACAACCCGTACGCTA |
| *IRX1* | CAAGAATCCCTACCCCACCA | TCCCCATGTCACCTTGTTCT |
| *IRX2* | TCACCAAGATGACCCTCACC | TTCGCTTTTGTTTCTCGGGG |
| *ISL1* | CTGTGGACATTACTCCCTCTTAC | GCAACCAACACATAGGGAAATC |
| *KCNJ11* | GCGCTTTGTGCCCATTGTA | TTGACGGTGTTGCCAAACTTG |
| *LMX1A* | TCCTAGCCTTGGAGAAGCAACT | CAGTGACTGGAGCAGAGAGAA |
| *LMX1B* | ACCTCCTTAACCAGCCTCAG | GCATGGAGTAGAGCCGGTC |
| *MAFB* | GGAGAATGAGAAGACGCAGC | GTTTCTCGCACTTGACCTTGT |
| *NEUROD1* | GCCCCAGGGTTATGAGACTAT | GAGAACTGAGACACTCGTCTGT |
| *NEUROG3* | GGCTGTGGGTGCTAAGGGTAAG | CAGGGAGAAGCAGAAGGAACAA |
| *NKX2.2* | AAACCATGTCACGCGCTCA | GGCGTTGTACTGCATGTGCT |
| *NKX6.1* | GGGCTCGTTTGGCCTATTCGTT | CCACTTGGTCCGGCGGTTCT |
| *ONECUT1* | GGACCTCAAGATAGCAGGTTTAT | CAGAATGCAGGTGAGCTAAGT |
| *ONECUT2* | GCCATCTTCAAGGAGAACAAAC | CGTTCATGAAGAAGTTGCTGAC |
| *PAX4* | AGCAGAGGCACTGGAGAAAGAGTT | CAGCTGCATTTCCCACTTGAGCTT |
| *PAX6* | GCGGAAGCTGCAAAGAAATAG | GGGCAAACACATCTGGATAATG |
| *PCSK1* | TCACACATGGGGAGAGAACC | TCCCGTGCAAAATCAGCTTC |
| *PCSK2* | CGGGTTCCTCTTCTGTGTCA | GACTCCAAAGCCGTGTTCTG |
| *PDX1* | CGTCCAGCTGCCTTTCCCAT | CCGTGAGATGTACTTGTTGAATAGGA |
| *PPY* | AGGTGCTCGCTTGGTCTAGTG | ACCCAGCAGTGGCTGTAGTAAC |
| *PTF1A* | CCAGAAGGTCATCATCTGCC | AGAGAGTGTCCTGCTAGGGG |
| *PTPRN2* | ACTGAGGATGTGGAGAAGGC | TGAGTTTGCTTTTCGACCCG |
| *RFX3* | AGGAACACAAACTGGACCCA | AGAGGGGAATCTGGCTTGAC |

| | | |
|---|---|---|
| *RFX3* genomic DNA | CAACAACAGGTACAGCAGGT | GATTCAGATGGGCGTCACAG |
| *RFX6* | GTCGATGCATGGCTTGGACT | TGGGCCATAGCTAGACGGTG |
| *SCG3* | GGGACTCCTTTAACCGCTGA | TCCAGTGTATGTGCTTGGCT |
| *SIX3* | GCGACTCGGAATGTGATGTAT | GGAGAAGGAAGAGGAGGAAGA |
| *SLC18A1* | ATGGTCATCACTGGGGTCAT | GGCTTCTGGGTTGCATACAT |
| *SLC30A8* | TGGGAGTCTTGCTGTTGTCA | CATCCAAATGTCAGCCGCTT |
| *SOX9* | GACTACACCGACCACCAGAACTCC | GTCTGCGGGATGGAAGGGA |
| *SST* | AGCTGCTGTCTGAACCCAAC | CCATAGCCGGGTTTGAGTTA |
| *TPH1* | CCCTTCTATACCCCAGAGCC | CCAAGAGAAGCCAAGCCAAT |
| *TXNIP* | GATACCCCAGAAGCTCCTCC | TGAACTTGAACTCAGGGGCA |
| *UCN3* | GATGGGCTTGGCTTTGTAGA | GGAGGGAAGTCCACTCTCG |

**Table S3.** List of antibodies used in the study.

| Antibody | Catalog number | Company | RRID | Dilution |
|---|---|---|---|---|
| Anti-β-Actin | Sc-47778 | Santa Cruz Biotechnology | AB_626632 | 1:10,000 (WB) |
| Anti-CDX2 | ab76541 | Abcam | AB_1523334 | 1:2000 (WB) |
| Anti-FEV | 25058-1-AP | Proteintech | AB_2879877 | 1:1000 (IF) |
| Anti-FOXA2 | 3143 | Cell Signaling Technology | AB_2104878 | 1:1000 (IF) |
| Anti-OCT4 | 4286S | Cell Signaling Technology | AB_1904076 | 1:500 (IF) |
| Anti-NANOG | 9656s | Cell Signaling Technology | AB_1658242 | 1:500 (IF) |
| Anti-SSEA4 | 9656s | Cell Signaling Technology | AB_1658242 | 1:500 (IF) |
| Anti-TRA-1-60 | 9656s | Cell Signaling Technology | AB_1658242 | 1:500 (IF) |
| Anti-SOX17 | AF1924 | R & D Systems | AB_355060 | 1:2000 (IF) |
| Anti-SOX9 | HPA001758 | Sigma | AB_1080067 | 1:1000 (IF), 1:4000 (WB) |
| Anti-GATA6 | AF1700 | R & D Systems | AB_2108901 | 1:1000 (IF) |
| Anti-PDX1 | ab47308 | Abcam | AB_777178 | 1:1000 (IF), 1:100 (FACS) |
| Anti-NKX6.1 | F55A12 | DSHB | AB_532379 | 1:2000 (IF), 1:100 (FACS) |
| Anti-NKX2.2 | 74.5A5-c | DSHB | AB_531794 | 1:2000 (IF) |
| Anti-NEUROG3 | AF3444 | R & D Systems | AB_2149527 | 1:1000 (IF) |
| Anti-RFX3 | NBP1-86301 | Novus Biologicals | AB_11019457 | 1:1000 (IF), 1:500 (WB) |
| Anti-CHROMOGRANIN A | MA5-14536 | Invitrogen | AB_10978165 | 1:2000 (IF) |
| Anti-TXNIP | ab215366 | Abcam | – | 1:2000 (WB) |
| Anti-INSULIN | GN-ID4-s | DSHB | AB_2255626 | 1:2000 (IF), 1:100 (FACS) |
| Anti-GLUCAGON | G2654 | Sigma | AB_259852 | 1:2000 (IF) |
| Anti-SOMATOSTATIN | MAB354 | Millipore | AB_2255365 | 1:2000 (IF) |
| Anti-UROCORTIN 3 | HPA038281 | Sigma | AB_10672408 | 1:1000 (IF) |
| Anti-PANCREATIC POLYPEPTIDE Y | ab113694 | Abcam | AB_11156699 | 1:1000 (IF) |
| Anti-GHRELIN | 5992 | BioVision | AB_2111582 | 1:1000 (IF) |
| Alexa Fluor 488 anti-rabbit IgG | A21206 | Invitrogen | AB_2535792 | 1:500 (IF, FACS) |
| Alexa Fluor 568 anti-rabbit IgG | A10042 | Invitrogen | AB_2534017 | 1:500 (IF) |

| | | | | |
|---|---|---|---|---|
| Alexa Fluor 488 anti-mouse IgG | A21202 | Invitrogen | AB_141607 | 1:500 (IF, FACS) |
| Alexa Fluor 594 anti-mouse IgG | A32744 | Invitrogen | AB_2762826 | 1:500 (IF) |
| Alexa Fluor 647 anti-mouse IgG | A31571 | Invitrogen | AB_162542 | 1:500 (FACS) |
| Alexa Fluor 488 anti-sheep IgG | A11015 | Invitrogen | AB_2534082 | 1:500 (IF) |
| Alexa Fluor 488 anti-guinea pig IgG | A11073 | Invitrogen | AB_2534117 | 1:500 (IF) |
| Alexa Fluor 647 anti-guinea pig IgG | A-21450 | Invitrogen | AB_2535867 | 1:500 (FACS) |
| Alexa Fluor 488 anti-goat IgG | A-11055 | Invitrogen | AB_2534102 | 1:500 (IF) |
| Alexa Fluor 568 anti-guinea pig IgG | A-11077 | Invitrogen | AB_2534121 | 1:500 (IF, FACS) |
| Peroxidase AffiniPure Donkey anti-Rabbit IgG (H+L) | 711-035-152 | Jackson ImmunoResearch Laboratories | AB_10015282 | 1:10,000 (WB) |
| Peroxidase AffiniPure Donkey anti-Mouse IgG (H+L) | 715-035-150 | Jackson ImmunoResearch Laboratories | AB_2340770 | 1:10,000 (WB) |
| Peroxidase-AffiniPure Donkey Anti-Guinea Pig IgG (H+L) | 706-035-148 | Jackson ImmunoResearch Laboratories | AB_2340447 | 1:10,000 (WB) |

**Table S4.** Top differentially expressed genes (DEGs) ($Log_2$ FC < -1, $p < 0.05$) downregulated in *RFX3* KO1 and *RFX3* KO2 at pancreatic progenitor (PP) stage compared to WT.

| Gene ID | $Log_2$ Fold Change | *p*-value |
|---|---|---|
| *GCG* | -9.432 | 0.0E+00 |
| *UCN3* | -8.572 | 1.6E-16 |
| *SST* | -6.520 | 1.0E-187 |
| *KCNK16* | -5.536 | 6.9E-61 |
| *INS* | -5.294 | 7.1E-83 |
| *IL20RA* | -5.211 | 1.2E-51 |
| *ARX* | -5.187 | 4.5E-61 |
| *GHRL* | -4.998 | 1.9E-95 |
| *CNGA3* | -4.907 | 1.6E-27 |
| *CRYBA2* | -4.876 | 7.6E-87 |
| *GPR119* | -4.786 | 1.5E-64 |
| *UCA1* | -4.734 | 3.8E-03 |
| *ABCC8* | -4.433 | 3.1E-36 |
| *VSTM2L* | -4.306 | 2.4E-47 |
| *ISL1* | -4.288 | 7.0E-69 |
| *CRH* | -4.210 | 1.6E-22 |
| *BRINP2* | -4.168 | 3.1E-23 |
| *ERICH3* | -4.063 | 1.9E-20 |
| *GAD2* | -3.990 | 1.8E-30 |
| *CACNA1A* | -3.976 | 2.9E-38 |
| *NSG2* | -3.965 | 6.0E-11 |
| *IRX1* | -3.886 | 2.0E-13 |
| *FGF14* | -3.870 | 1.0E-30 |
| *MAFB* | -3.823 | 1.2E-64 |
| *PSCA* | -3.790 | 7.8E-10 |
| *GJD2* | -3.745 | 6.9E-10 |
| *ATP2A3* | -3.739 | 1.3E-73 |
| *KCNH6* | -3.651 | 7.2E-44 |
| *PERCC1* | -3.615 | 1.2E-20 |
| *ADGRA1* | -3.578 | 2.3E-14 |
| *PAX6* | -3.532 | 5.9E-16 |
| *LHFPL4* | -3.521 | 2.5E-33 |
| *PTPRN* | -3.492 | 6.9E-51 |
| *TRPM3* | -3.479 | 2.7E-21 |

| | | |
|---|---|---|
| *SEZ6L* | -3.478 | 4.7E-84 |
| *LY6H* | -3.442 | 2.5E-10 |
| *RGS6* | -3.408 | 1.8E-20 |
| *TNR* | -3.355 | 1.4E-16 |
| *C11orf87* | -3.305 | 9.7E-23 |
| *SSTR2* | -3.228 | 2.3E-54 |
| *PTPRN2* | -3.194 | 4.3E-86 |
| *NECAB2* | -3.189 | 7.3E-18 |
| *SYT5* | -3.169 | 1.6E-10 |
| *NEGR1* | -3.104 | 9.6E-16 |
| *GCK* | -3.051 | 5.0E-17 |
| *ST6GALNAC5* | -3.018 | 1.9E-21 |
| *ASCL2* | -2.978 | 3.3E-10 |
| *SMIM32* | -2.974 | 2.4E-18 |
| *RYR1* | -2.964 | 5.7E-20 |
| *UNC5A* | -2.950 | 1.7E-19 |
| *PCSK1N* | -2.892 | 6.2E-23 |
| *ASCL1* | -2.890 | 1.6E-10 |
| *TMEM196* | -2.852 | 4.1E-20 |
| *THSD4* | -2.832 | 5.6E-54 |
| *SGCD* | -2.823 | 9.3E-24 |
| *KCNK17* | -2.808 | 1.9E-12 |
| *GRIN3A* | -2.790 | 9.4E-28 |
| *GC* | -2.782 | 8.4E-28 |
| *AQP10* | -2.780 | 3.3E-15 |
| *FFAR2* | -2.777 | 2.5E-15 |
| *TMEM130* | -2.765 | 2.2E-08 |
| *CHST8* | -2.761 | 2.0E-16 |
| *CFC1* | -2.753 | 1.3E-07 |
| *GDAP1L1* | -2.744 | 1.0E-15 |
| *PKHD1L1* | -2.718 | 1.6E-12 |
| *KIF19* | -2.711 | 4.9E-17 |
| *UNC13A* | -2.710 | 9.9E-21 |
| *USH2A* | -2.700 | 2.6E-16 |
| *GPBAR1* | -2.696 | 1.3E-15 |
| *KCND3* | -2.679 | 4.0E-42 |
| *FSTL5* | -2.655 | 3.4E-36 |
| *CAMK2B* | -2.648 | 7.1E-25 |

| | | |
|---|---|---|
| *MARCHF4* | -2.603 | 1.5E-09 |
| *SCG3* | -2.599 | 8.2E-20 |
| *SYP* | -2.599 | 1.9E-17 |
| *GRIK1* | -2.596 | 4.7E-10 |
| *CHGB* | -2.580 | 1.1E-24 |
| *PDE2A* | -2.578 | 1.7E-17 |
| *RIMBP2* | -2.568 | 4.2E-44 |
| *BAIAP3* | -2.565 | 5.7E-57 |
| *LRRC10B* | -2.548 | 1.2E-10 |
| *CLDN18* | -2.539 | 3.1E-56 |
| *GNAO1* | -2.538 | 4.0E-21 |
| *ERO1B* | -2.535 | 1.0E-44 |
| *CALY* | -2.522 | 8.8E-20 |
| *ST18* | -2.516 | 2.8E-15 |
| *SLC7A14* | -2.506 | 4.3E-16 |
| *VWA5B2* | -2.491 | 2.5E-56 |
| *GNG2* | -2.487 | 5.8E-17 |
| *GDPD2* | -2.485 | 2.7E-11 |
| *CELF3* | -2.479 | 2.4E-48 |
| *CDK5R2* | -2.466 | 2.8E-16 |
| *ASIC4* | -2.463 | 1.3E-10 |
| *NEUROD1* | -2.455 | 7.9E-34 |
| *RASGRF2* | -2.455 | 2.2E-22 |
| *STUM* | -2.446 | 1.4E-11 |
| *KCNB1* | -2.434 | 3.7E-11 |
| *PLCXD3* | -2.361 | 6.4E-13 |
| *PCSK1* | -2.345 | 2.6E-30 |
| *TENM2* | -2.328 | 8.3E-12 |
| *FBXL16* | -2.312 | 8.0E-22 |
| *GRAMD2A* | -2.311 | 7.4E-08 |
| *AMER3* | -2.291 | 1.3E-13 |
| *KCNJ6* | -2.287 | 4.5E-20 |
| *C22orf42* | -2.285 | 5.6E-13 |
| *KCNH3* | -2.278 | 1.2E-24 |
| *RAB26* | -2.274 | 3.6E-74 |
| *RUNDC3A* | -2.272 | 1.2E-22 |
| *SPTBN4* | -2.253 | 6.7E-08 |
| *POU2F2* | -2.235 | 1.9E-11 |

| | | |
|---|---|---|
| *UNC80* | -2.231 | 4.5E-27 |
| *CHGA* | -2.230 | 8.7E-19 |
| *DPEP1* | -2.229 | 9.0E-16 |
| *DISP2* | -2.207 | 1.7E-27 |
| *ELAVL4* | -2.201 | 2.0E-09 |
| *IRX2* | -2.197 | 2.0E-14 |
| *ANKS1B* | -2.171 | 1.5E-24 |
| *KCNMA1* | -2.169 | 3.2E-10 |
| *C1QL1* | -2.163 | 1.7E-21 |
| *PGM5* | -2.158 | 6.9E-30 |
| *AMPH* | -2.158 | 1.6E-14 |
| *MGAM2* | -2.157 | 2.0E-15 |
| *TUNAR* | -2.150 | 4.5E-25 |
| *CPLX2* | -2.142 | 9.9E-08 |
| *TSPAN1* | -2.118 | 1.2E-09 |
| *FEV* | -2.111 | 1.6E-18 |
| *NRXN1* | -2.093 | 5.1E-12 |
| *UBE2QL1* | -2.092 | 3.2E-09 |
| *RUNX1T1* | -2.084 | 4.4E-16 |
| *CHRNB2* | -2.069 | 9.5E-12 |
| *AK5* | -2.062 | 8.0E-21 |
| *NFASC* | -2.051 | 3.1E-30 |
| *KCNJ5* | -2.048 | 1.4E-06 |
| *ATCAY* | -2.046 | 4.0E-14 |
| *RAPGEF4* | -2.038 | 7.9E-24 |
| *SVOP* | -2.030 | 3.2E-08 |
| *DLL4* | -2.024 | 1.1E-08 |
| *SYT4* | -2.014 | 3.1E-14 |
| *ADAM12* | -1.996 | 3.0E-09 |
| *CORO2B* | -1.985 | 1.3E-11 |
| *ADCY2* | -1.979 | 6.8E-20 |
| *NEUROG3* | -1.972 | 9.2E-21 |
| *CTNNA2* | -1.968 | 1.6E-18 |
| *NKX2-2* | -1.960 | 2.6E-36 |
| *SPOCK3* | -1.957 | 1.2E-15 |
| *SLIT1* | -1.952 | 3.1E-18 |
| *SLC8A1* | -1.952 | 1.1E-06 |
| *RAP1GAP2* | -1.940 | 1.9E-30 |

| | | |
|---|---|---|
| *INSM1* | -1.939 | 7.1E-21 |
| *MMRN1* | -1.935 | 2.3E-04 |
| *ADAMTS2* | -1.931 | 4.2E-09 |
| *SSTR1* | -1.922 | 2.0E-20 |
| *MAP6* | -1.920 | 3.4E-37 |
| *SLC6A4* | -1.920 | 1.0E-13 |
| *SPOCK1* | -1.911 | 1.1E-04 |
| *SLC5A9* | -1.900 | 2.2E-15 |
| *CNIH2* | -1.899 | 9.5E-28 |
| *DEPP1* | -1.891 | 2.5E-12 |
| *PNMA2* | -1.891 | 9.4E-21 |
| *SYT7* | -1.882 | 1.3E-21 |
| *MANEAL* | -1.878 | 3.3E-08 |
| *LMX1B* | -1.873 | 2.0E-59 |
| *CELA3A* | -1.868 | 2.8E-09 |
| *RAB3C* | -1.866 | 5.3E-18 |
| *DUSP26* | -1.864 | 6.0E-12 |
| *DDC* | -1.864 | 8.3E-31 |
| *SEC14L5* | -1.863 | 9.8E-06 |
| *MSC* | -1.858 | 1.2E-12 |
| *NPY5R* | -1.857 | 1.9E-04 |
| *LGALS12* | -1.849 | 2.3E-10 |
| *CACNA1C* | -1.840 | 1.3E-17 |
| *ELAPOR1* | -1.829 | 5.6E-40 |
| *STMN2* | -1.821 | 3.4E-11 |
| *CDHR3* | -1.821 | 2.7E-13 |
| *HPSE* | -1.810 | 8.1E-25 |
| *CKMT1B* | -1.803 | 8.6E-06 |
| *LMO3* | -1.800 | 1.7E-20 |
| *DISP3* | -1.775 | 1.7E-13 |
| *CACNA1B* | -1.775 | 3.9E-17 |
| *MAPK11* | -1.768 | 1.9E-14 |
| *PNPLA7* | -1.764 | 5.0E-12 |
| *LINGO2* | -1.756 | 1.3E-15 |
| *RSAD2* | -1.756 | 1.0E-08 |
| *ADAMTS5* | -1.753 | 1.2E-08 |
| *HEPACAM2* | -1.746 | 3.5E-20 |
| *FAM181B* | -1.709 | 3.2E-11 |

| | | |
|---|---|---|
| *TPPP3* | -1.701 | 4.3E-09 |
| *DRD2* | -1.700 | 3.8E-04 |
| *CPA1* | -1.698 | 3.4E-13 |
| *GALR1* | -1.686 | 8.4E-12 |
| *TRIM50* | -1.683 | 2.4E-07 |
| *C7* | -1.669 | 3.6E-05 |
| *SLC26A3* | -1.653 | 4.2E-04 |
| *GIPR* | -1.641 | 8.7E-18 |
| *AP3B2* | -1.641 | 4.8E-36 |
| *HS3ST4* | -1.635 | 4.2E-12 |
| *UNC79* | -1.634 | 5.0E-18 |
| *CD177* | -1.627 | 2.3E-11 |
| *IGSF10* | -1.623 | 9.3E-15 |
| *CGA* | -1.619 | 7.6E-07 |
| *SLC35G2* | -1.618 | 7.8E-10 |
| *PCSK2* | -1.613 | 3.1E-05 |
| *PPP1R1A* | -1.609 | 4.4E-13 |
| *SNAP91* | -1.608 | 2.1E-11 |
| *GATM* | -1.608 | 3.9E-18 |
| *RGS9* | -1.606 | 1.0E-13 |
| *CNNM1* | -1.604 | 5.7E-19 |
| *NYAP1* | -1.586 | 7.1E-10 |
| *GFRA3* | -1.585 | 9.9E-13 |
| *SCN3A* | -1.584 | 1.0E-08 |
| *DPP6* | -1.581 | 2.0E-13 |
| *DNAH5* | -1.570 | 6.9E-06 |
| *SCGN* | -1.565 | 7.4E-28 |
| *NPHS1* | -1.565 | 1.3E-12 |
| *MAMLD1* | -1.541 | 1.2E-26 |
| *ENPP1* | -1.541 | 4.4E-08 |
| *LRRTM1* | -1.539 | 1.4E-04 |
| *PLXNA2* | -1.536 | 6.9E-23 |
| *ACVR1C* | -1.535 | 1.3E-10 |
| *PLD5* | -1.534 | 5.8E-04 |
| *CTSE* | -1.532 | 2.2E-05 |
| *MYOCD* | -1.529 | 3.4E-03 |
| *BSN* | -1.522 | 2.1E-20 |
| *CNTNAP4* | -1.522 | 6.0E-08 |

| | | |
|---|---|---|
| *TMOD1* | -1.520 | 4.0E-20 |
| *SNED1* | -1.514 | 1.6E-10 |
| *CDH12* | -1.514 | 8.7E-09 |
| *SV2B* | -1.511 | 1.2E-06 |
| *RGS4* | -1.509 | 1.7E-07 |
| *SSTR3* | -1.499 | 6.1E-05 |
| *FUT2* | -1.497 | 1.5E-07 |
| *CRB2* | -1.496 | 1.0E-12 |
| *IGFBPL1* | -1.495 | 3.0E-07 |
| *SLC25A53* | -1.491 | 7.2E-17 |
| *SLC38A3* | -1.488 | 1.5E-66 |
| *PDZD2* | -1.485 | 4.1E-13 |
| *PRUNE2* | -1.480 | 1.6E-18 |
| *ASTN1* | -1.475 | 1.5E-03 |
| *BAALC* | -1.467 | 4.8E-13 |
| *GRIA2* | -1.467 | 5.2E-06 |
| *KIF5A* | -1.466 | 8.7E-16 |
| *MLXIPL* | -1.461 | 7.2E-14 |
| *KCNK3* | -1.460 | 9.2E-08 |
| *PSD* | -1.459 | 9.5E-11 |
| *PLPPR1* | -1.452 | 1.8E-05 |
| *NEURL1* | -1.450 | 1.3E-14 |
| *LGI2* | -1.443 | 1.1E-08 |
| *LRP1B* | -1.440 | 2.0E-07 |
| *NOVA2* | -1.435 | 6.4E-04 |
| *NOL4* | -1.434 | 1.5E-08 |
| *GP2* | -1.424 | 7.7E-09 |
| *SMPD3* | -1.424 | 1.1E-07 |
| *ZNF506* | -1.420 | 2.9E-17 |
| *MYPN* | -1.419 | 1.3E-10 |
| *XKR7* | -1.417 | 4.1E-09 |
| *MAPK8IP2* | -1.412 | 1.4E-19 |
| *RASA4B* | -1.406 | 4.0E-16 |
| *ADAMTSL1* | -1.406 | 3.1E-11 |
| *KCNH2* | -1.405 | 2.4E-19 |
| *ADGRF3* | -1.388 | 4.6E-06 |
| *OPRK1* | -1.387 | 7.9E-03 |
| *PRKCG* | -1.387 | 7.0E-05 |

| | | |
|---|---|---|
| *FMN2* | -1.384 | 3.8E-08 |
| *SCG2* | -1.380 | 2.8E-07 |
| *FRRS1L* | -1.376 | 4.6E-17 |
| *CADM2* | -1.372 | 8.1E-07 |
| *RHOH* | -1.370 | 8.7E-05 |
| *ST8SIA5* | -1.368 | 2.9E-03 |
| *HEYL* | -1.368 | 1.0E-08 |
| *VLDLR* | -1.365 | 1.6E-07 |
| *GRIK3* | -1.353 | 1.7E-02 |
| *SLC25A12* | -1.353 | 8.9E-12 |
| *GRHL3* | -1.349 | 4.3E-11 |
| *NAV3* | -1.347 | 2.7E-06 |
| *SMIM5* | -1.339 | 3.5E-04 |
| *TUBA4A* | -1.332 | 5.7E-19 |
| *DIRAS3* | -1.328 | 1.7E-03 |
| *RTL9* | -1.327 | 1.6E-07 |
| *STX1A* | -1.327 | 2.4E-25 |
| *RIMS2* | -1.326 | 1.7E-13 |
| *NPR1* | -1.324 | 7.8E-03 |
| *KIAA0319* | -1.323 | 2.4E-05 |
| *TEKT2* | -1.323 | 3.8E-13 |
| *AMHR2* | -1.314 | 2.6E-17 |
| *BTBD11* | -1.313 | 2.1E-08 |
| *CPA4* | -1.312 | 2.4E-07 |
| *QPCT* | -1.311 | 8.1E-15 |
| *SLC24A3* | -1.311 | 2.5E-02 |
| *SPAG6* | -1.305 | 2.7E-07 |
| *MNX1* | -1.300 | 1.9E-21 |
| *CLMN* | -1.291 | 2.7E-26 |
| *TACR1* | -1.287 | 4.7E-05 |
| *GAL* | -1.286 | 4.6E-04 |
| *TAGLN3* | -1.285 | 4.5E-08 |
| *ADORA2A* | -1.279 | 1.1E-26 |
| *RASA4* | -1.277 | 2.4E-05 |
| *ACSL1* | -1.272 | 1.1E-29 |
| *IL32* | -1.268 | 2.7E-05 |
| *GFRA1* | -1.263 | 2.8E-06 |
| *ZFPM2* | -1.256 | 4.2E-14 |

| | | |
|---|---|---|
| *TTLL6* | -1.249 | 2.1E-05 |
| *NANOS1* | -1.247 | 1.6E-11 |
| *KIRREL2* | -1.244 | 1.2E-04 |
| *GABRB3* | -1.243 | 1.3E-18 |
| *PLAT* | -1.239 | 3.2E-02 |
| *VGF* | -1.234 | 2.5E-05 |
| *HTRA1* | -1.229 | 5.1E-04 |
| *CELF4* | -1.229 | 1.8E-12 |
| *RTL1* | -1.225 | 6.6E-05 |
| *CASZ1* | -1.224 | 2.6E-07 |
| *LSAMP* | -1.221 | 1.4E-09 |
| *ESM1* | -1.220 | 3.8E-02 |
| *FNDC11* | -1.215 | 1.9E-03 |
| *STC2* | -1.212 | 4.6E-14 |
| *SLC16A12* | -1.208 | 2.6E-06 |
| *RDH12* | -1.206 | 7.1E-08 |
| *POMC* | -1.204 | 8.4E-04 |
| *SLC2A14* | -1.202 | 2.4E-15 |
| *XKR4* | -1.199 | 6.8E-11 |
| *PPP1R3C* | -1.187 | 6.1E-10 |
| *SRRM4* | -1.177 | 5.6E-09 |
| *RASGRP1* | -1.170 | 1.6E-07 |
| *TM4SF4* | -1.169 | 8.3E-07 |
| *HYDIN* | -1.168 | 1.9E-10 |
| *ZDHHC11B* | -1.164 | 4.6E-10 |
| *ATP8A1* | -1.160 | 2.2E-08 |
| *TMEM132B* | -1.158 | 3.5E-04 |
| *PTF1A* | -1.156 | 2.1E-04 |
| *KCNJ11* | -1.152 | 3.5E-05 |
| *RASSF6* | -1.148 | 2.5E-08 |
| *RFX6* | -1.144 | 2.5E-13 |
| *TTBK1* | -1.139 | 2.6E-10 |
| *FAM167A* | -1.136 | 1.8E-06 |
| *CADPS* | -1.135 | 2.3E-10 |
| *CAMK1D* | -1.134 | 9.6E-08 |
| *PDE1C* | -1.133 | 7.3E-05 |
| *SBK2* | -1.131 | 2.8E-02 |
| *TLE6* | -1.127 | 4.8E-09 |

| | | |
|---|---|---|
| *KCNV1* | -1.123 | 2.6E-03 |
| *KCNQ2* | -1.122 | 2.5E-03 |
| *SCN5A* | -1.121 | 2.9E-07 |
| *OSGIN1* | -1.121 | 6.6E-05 |
| *LRRC24* | -1.117 | 9.4E-04 |
| *CPA2* | -1.116 | 1.1E-04 |
| *GNG4* | -1.107 | 2.8E-10 |
| *ACACB* | -1.106 | 5.0E-10 |
| *ANGPT2* | -1.102 | 3.0E-03 |
| *ERN1* | -1.101 | 7.5E-09 |
| *NBEAL2* | -1.098 | 3.0E-35 |
| *EGLN3* | -1.094 | 4.3E-04 |
| *UBD* | -1.090 | 1.7E-03 |
| *PLXNC1* | -1.089 | 5.1E-21 |
| *WNT9A* | -1.089 | 1.6E-05 |
| *CARMIL3* | -1.086 | 3.8E-05 |
| *WDR17* | -1.081 | 1.0E-05 |
| *ECE1* | -1.079 | 2.5E-09 |
| *C2CD2L* | -1.079 | 2.3E-06 |
| *SMARCA2* | -1.072 | 1.8E-12 |
| *STXBP5L* | -1.072 | 1.0E-08 |
| *COPG2IT1* | -1.071 | 4.3E-07 |
| *PDK3* | -1.069 | 3.5E-10 |
| *STAC* | -1.068 | 1.4E-03 |
| *C4A* | -1.062 | 1.4E-07 |
| *HSFX2* | -1.059 | 6.0E-05 |
| *CD82* | -1.057 | 2.7E-06 |
| *H2BC21* | -1.054 | 5.5E-06 |
| *APC2* | -1.053 | 1.1E-04 |
| *MARCHF8* | -1.051 | 2.5E-50 |
| *ELMO1* | -1.046 | 3.6E-06 |
| *MAPRE3* | -1.044 | 4.8E-21 |
| *ERP27* | -1.043 | 6.5E-13 |
| *RIMKLA* | -1.040 | 4.9E-08 |
| *TACSTD2* | -1.038 | 2.0E-02 |
| *GCNT1* | -1.034 | 9.3E-10 |
| *SPATA13* | -1.034 | 6.6E-24 |
| *PPFIA3* | -1.034 | 1.9E-07 |

| | | |
|---|---|---|
| *ALDH1A1* | -1.027 | 3.0E-07 |
| *CKMT1A* | -1.027 | 1.9E-03 |
| *CACNA1I* | -1.025 | 3.4E-04 |
| *SEZ6* | -1.025 | 1.3E-02 |
| *CKMT2* | -1.024 | 3.8E-07 |
| *SNAP25* | -1.023 | 7.7E-10 |
| *MAPK15* | -1.023 | 8.2E-04 |
| *B3GALT4* | -1.020 | 3.7E-04 |
| *GRIK2* | -1.007 | 5.7E-05 |
| *SOX1* | -1.006 | 3.1E-05 |
| *SPOCK2* | -1.000 | 1.2E-03 |

**Table S5.** Top differentially expressed genes (DEGs) ($Log_2$ FC > 1, $p < 0.05$) upregulated in *RFX3* KO1 and *RFX3* KO2 at pancreatic progenitor (PP) stage compared to WT.

| **Gene ID** | **Log2 Fold Change** | ***p*-value** |
|---|---|---|
| *APOA4* | 2.695 | 1.1E-22 |
| *ITIH3* | 2.585 | 3.2E-06 |
| *SLC22A7* | 2.459 | 1.5E-04 |
| *HRG* | 2.447 | 7.3E-11 |
| *ALDOB* | 2.359 | 5.1E-13 |
| *FGF18* | 2.252 | 4.5E-03 |
| *KNG1* | 2.131 | 2.0E-10 |
| *MAT1A* | 2.128 | 1.4E-19 |
| *ALB* | 2.030 | 2.3E-27 |
| *IP6K3* | 2.025 | 6.8E-03 |
| *RBP2* | 1.945 | 9.6E-08 |
| *SI* | 1.897 | 1.5E-25 |
| *ITIH1* | 1.892 | 1.3E-05 |
| *SPP1* | 1.889 | 2.8E-05 |
| *SERPINA7* | 1.823 | 1.2E-16 |
| *LRRK2* | 1.737 | 5.9E-17 |
| *HMGCS2* | 1.710 | 5.6E-07 |
| *TPH1* | 1.686 | 3.7E-04 |
| *PCAT1* | 1.637 | 3.1E-04 |
| *FGG* | 1.621 | 1.9E-39 |
| *UGT2B10* | 1.615 | 1.1E-07 |
| *FAM151A* | 1.610 | 1.3E-14 |
| *APOC2* | 1.602 | 2.8E-07 |
| *URAD* | 1.585 | 2.9E-04 |
| *CRLF1* | 1.582 | 2.6E-02 |
| *FGA* | 1.568 | 4.8E-30 |
| *FETUB* | 1.564 | 4.3E-04 |
| *F5* | 1.533 | 1.1E-02 |
| *ANGPTL3* | 1.519 | 1.2E-04 |
| *GUCY2C* | 1.509 | 1.3E-09 |
| *TDO2* | 1.496 | 8.1E-18 |
| *APOC3* | 1.496 | 4.9E-07 |
| *METTL7A* | 1.492 | 1.7E-05 |
| *ITIH2* | 1.483 | 1.5E-05 |
| *MMP1* | 1.481 | 8.3E-08 |

| | | |
|---|---|---|
| *CALB1* | 1.458 | 2.3E-07 |
| *TF* | 1.452 | 1.4E-10 |
| *CREB3L3* | 1.442 | 3.1E-11 |
| *SMLR1* | 1.436 | 1.4E-17 |
| *HRH2* | 1.436 | 2.7E-10 |
| *ANGPT1* | 1.434 | 1.2E-12 |
| *PCDHA9* | 1.431 | 2.2E-02 |
| *RFTN2* | 1.420 | 1.1E-06 |
| *ACSL5* | 1.400 | 1.5E-14 |
| *AKR1D1* | 1.385 | 3.7E-28 |
| *KCNJ13* | 1.379 | 2.0E-07 |
| *GSTA2* | 1.377 | 1.6E-45 |
| *PIK3C2G* | 1.369 | 5.8E-09 |
| *HPX* | 1.364 | 2.7E-12 |
| *C20orf204* | 1.357 | 4.8E-06 |
| *COL25A1* | 1.354 | 8.3E-04 |
| *SLITRK3* | 1.348 | 1.3E-07 |
| *AHSG* | 1.334 | 8.7E-08 |
| *HSD17B2* | 1.324 | 1.9E-11 |
| *ECM2* | 1.310 | 6.3E-07 |
| *PLG* | 1.307 | 3.7E-06 |
| *CLDN2* | 1.307 | 9.5E-06 |
| *GSTA1* | 1.287 | 9.9E-16 |
| *MEP1A* | 1.284 | 1.7E-18 |
| *MGAM* | 1.281 | 2.0E-05 |
| *KLB* | 1.280 | 1.3E-05 |
| *GPM6A* | 1.278 | 3.9E-04 |
| *SERPINA6* | 1.277 | 2.0E-04 |
| *APOB* | 1.271 | 2.8E-08 |
| *GBA3* | 1.260 | 3.7E-06 |
| *ARHGAP23* | 1.257 | 3.0E-05 |
| *PTCHD4* | 1.256 | 3.6E-03 |
| *DPYS* | 1.255 | 5.2E-13 |
| *L1TD1* | 1.248 | 1.4E-18 |
| *FGB* | 1.247 | 8.0E-29 |
| *CUBN* | 1.247 | 3.0E-19 |
| *SCHIP1* | 1.241 | 2.0E-03 |
| *COL11A1* | 1.240 | 1.1E-06 |

| | | |
|---|---|---|
| *FMO1* | 1.240 | 7.2E-07 |
| *CFTR* | 1.217 | 1.0E-10 |
| *AFP* | 1.198 | 9.3E-19 |
| *EGFLAM* | 1.188 | 5.2E-11 |
| *FABP1* | 1.186 | 1.3E-05 |
| *GDNF* | 1.182 | 3.7E-04 |
| *NRGN* | 1.179 | 2.2E-11 |
| *PRAP1* | 1.168 | 6.5E-04 |
| *SP8* | 1.168 | 8.6E-06 |
| *PALMD* | 1.166 | 1.2E-03 |
| *ALDH1L1* | 1.166 | 2.3E-06 |
| *UGT2B7* | 1.153 | 6.6E-07 |
| *CA2* | 1.144 | 1.9E-23 |
| *XYLB* | 1.140 | 9.1E-06 |
| *TFF1* | 1.135 | 1.3E-02 |
| *HOGA1* | 1.134 | 1.5E-05 |
| *SERPINC1* | 1.129 | 4.8E-03 |
| *MAOB* | 1.125 | 3.1E-06 |
| *A2M* | 1.120 | 3.6E-05 |
| *AJAP1* | 1.119 | 4.5E-02 |
| *LEAP2* | 1.118 | 1.7E-03 |
| *EGF* | 1.117 | 5.2E-03 |
| *FMO5* | 1.106 | 2.9E-04 |
| *IYD* | 1.100 | 2.2E-03 |
| *LIPC* | 1.097 | 1.0E-07 |
| *PCDHA10* | 1.094 | 1.6E-06 |
| *METTL7B* | 1.092 | 4.9E-06 |
| *RBP4* | 1.088 | 3.2E-11 |
| *SLC16A8* | 1.080 | 2.9E-02 |
| *ITIH4* | 1.072 | 3.4E-05 |
| *SOAT2* | 1.070 | 1.9E-08 |
| *PLP1* | 1.066 | 6.6E-03 |
| *MYB* | 1.063 | 2.1E-02 |
| *SLC13A5* | 1.058 | 2.6E-04 |
| *NDRG1* | 1.057 | 1.5E-06 |
| *CBSL* | 1.056 | 4.0E-02 |
| *GPR3* | 1.055 | 3.0E-02 |
| *WNT11* | 1.054 | 1.2E-05 |

| | | |
|---|---|---|
| *MAMDC2* | 1.053 | 7.7E-03 |
| *ADAMTS3* | 1.046 | 6.6E-03 |
| *INPP5D* | 1.039 | 1.7E-03 |
| *SLITRK6* | 1.032 | 3.2E-03 |
| *TNFRSF10C* | 1.026 | 5.6E-04 |
| *VEPH1* | 1.019 | 8.4E-05 |
| *TMEM59L* | 1.019 | 1.6E-02 |
| *RELN* | 1.018 | 2.3E-04 |
| *KYNU* | 1.014 | 2.0E-09 |
| *AGT* | 1.008 | 2.9E-08 |
| *KLK6* | 1.006 | 7.8E-05 |
| *ADAMTS12* | 1.003 | 9.9E-10 |
| *DGAT2* | 1.002 | 1.8E-05 |
| *FAS* | 1.002 | 1.4E-02 |

**Table S6.** Top differentially expressed genes (DEGs) ($Log_2$ FC < -1, $p < 0.05$) downregulated in *RFX3* KO1 and *RFX3* KO2 at islet stage (S6) compared to WT.

| Gene ID | $Log_2$ Fold Change | *p*-value |
|---|---|---|
| *GCG* | -6.593 | 1.5E-106 |
| *IBSP* | -5.126 | 2.0E-15 |
| *SLC30A8* | -4.957 | 2.1E-36 |
| *LRRC53* | -4.763 | 4.5E-29 |
| *IAPP* | -4.763 | 1.8E-22 |
| *INS* | -4.611 | 9.6E-13 |
| *FFAR1* | -4.565 | 2.0E-07 |
| *SST* | -4.323 | 1.4E-48 |
| *PRG4* | -3.775 | 2.6E-32 |
| *SERINC4* | -3.554 | 3.7E-03 |
| *S100B* | -3.448 | 5.1E-08 |
| *BHMT* | -3.257 | 2.4E-05 |
| *KCNK16* | -3.183 | 9.9E-26 |
| *PAX5* | -3.123 | 2.4E-03 |
| *SLC6A17* | -3.121 | 1.1E-16 |
| *PAX6* | -2.868 | 5.1E-15 |
| *FGF17* | -2.861 | 7.3E-10 |
| *FEZF1* | -2.856 | 4.7E-05 |
| *GCK* | -2.798 | 3.7E-12 |
| *ANGPTL1* | -2.779 | 1.1E-03 |
| *CNGA3* | -2.751 | 2.9E-30 |
| *CDH8* | -2.713 | 1.3E-11 |
| *FGF1* | -2.710 | 1.6E-08 |
| *GAP43* | -2.673 | 1.6E-08 |
| *RMST* | -2.670 | 2.4E-22 |
| *MCHR1* | -2.656 | 1.8E-14 |
| *NR2E1* | -2.619 | 1.2E-02 |
| *FAP* | -2.610 | 7.8E-14 |
| *FEZF2* | -2.594 | 1.8E-03 |
| *ABCC8* | -2.586 | 2.2E-66 |
| *RGS1* | -2.580 | 7.3E-11 |
| *CACNA1E* | -2.557 | 2.4E-06 |
| *FABP7* | -2.550 | 1.2E-09 |
| *SIX6* | -2.544 | 4.4E-08 |
| *NPTX2* | -2.530 | 3.3E-08 |

| | | |
|---|---|---|
| *WNT8B* | -2.511 | 5.6E-04 |
| *LHX2* | -2.511 | 4.9E-04 |
| *SPARCL1* | -2.493 | 3.6E-09 |
| *NTNG2* | -2.480 | 1.1E-07 |
| *EGR3* | -2.453 | 4.6E-05 |
| *PLP1* | -2.424 | 4.1E-05 |
| *KIRREL3* | -2.409 | 1.0E-07 |
| *MEGF10* | -2.403 | 3.4E-05 |
| *TUBB1* | -2.397 | 2.4E-15 |
| *ISL1* | -2.385 | 4.9E-13 |
| *ERO1B* | -2.364 | 9.6E-37 |
| *SYT12* | -2.334 | 1.7E-06 |
| *CFI* | -2.325 | 3.2E-19 |
| *ZMYND10* | -2.311 | 7.8E-08 |
| *TMEM158* | -2.307 | 5.6E-09 |
| *FAM181A* | -2.286 | 5.7E-03 |
| *GABRQ* | -2.279 | 1.8E-06 |
| *WDR49* | -2.277 | 3.4E-03 |
| *SCN1A* | -2.268 | 3.4E-06 |
| *DLX1* | -2.264 | 3.7E-07 |
| *CRB1* | -2.260 | 1.1E-04 |
| *TPPP3* | -2.248 | 4.0E-04 |
| *DOCK10* | -2.240 | 1.1E-11 |
| *BTBD17* | -2.229 | 1.8E-05 |
| *ZIC2* | -2.225 | 9.2E-06 |
| *LMO1* | -2.225 | 5.4E-05 |
| *SUCNR1* | -2.218 | 3.6E-08 |
| *RTL1* | -2.211 | 3.6E-07 |
| *NTN1* | -2.207 | 1.1E-04 |
| *AMER2* | -2.207 | 7.6E-04 |
| *GAD2* | -2.203 | 1.4E-57 |
| *ARX* | -2.201 | 3.5E-05 |
| *LPL* | -2.182 | 1.5E-08 |
| *F13A1* | -2.181 | 2.0E-03 |
| *CORIN* | -2.179 | 4.7E-06 |
| *C6orf118* | -2.175 | 5.0E-03 |
| *ZIC3* | -2.161 | 6.1E-04 |
| *SHC3* | -2.153 | 2.4E-19 |

| | | |
|---|---|---|
| *VCAM1* | -2.150 | 4.2E-06 |
| *CYP26C1* | -2.142 | 9.7E-05 |
| *RYR1* | -2.140 | 1.3E-08 |
| *CALB2* | -2.131 | 2.5E-11 |
| *GRIK4* | -2.130 | 2.6E-06 |
| *NEUROG3* | -2.128 | 1.0E-06 |
| *PAX2* | -2.120 | 3.2E-02 |
| *ERICH3* | -2.109 | 5.0E-14 |
| *PTPRO* | -2.107 | 5.7E-07 |
| *PTPRZ1* | -2.105 | 4.4E-07 |
| *GPM6B* | -2.087 | 3.2E-08 |
| *CRIP2* | -2.079 | 4.8E-15 |
| *ZIC1* | -2.073 | 4.1E-05 |
| *EGR2* | -2.065 | 2.8E-12 |
| *SALL3* | -2.053 | 2.9E-07 |
| *NKX6-2* | -2.050 | 7.6E-12 |
| *PDE1A* | -2.047 | 1.1E-05 |
| *FOXG1* | -2.043 | 1.0E-03 |
| *EPHA5* | -2.032 | 4.4E-06 |
| *USH2A* | -2.031 | 5.8E-08 |
| *LY6H* | -2.012 | 1.1E-02 |
| *ZIC5* | -2.008 | 2.4E-03 |
| *ADD2* | -1.997 | 6.0E-04 |
| *APCDD1* | -1.994 | 1.2E-11 |
| *MAPK15* | -1.985 | 1.2E-31 |
| *SLC17A8* | -1.973 | 2.2E-16 |
| *SLC1A2* | -1.968 | 8.1E-03 |
| *NSG2* | -1.967 | 8.8E-03 |
| *DPP6* | -1.962 | 3.6E-10 |
| *ZDHHC22* | -1.960 | 3.5E-02 |
| *NTRK2* | -1.951 | 1.9E-12 |
| *SPAG6* | -1.951 | 4.4E-12 |
| *KCNK17* | -1.950 | 1.5E-08 |
| *MLC1* | -1.941 | 1.4E-04 |
| *HYDIN* | -1.927 | 8.8E-08 |
| *GRPR* | -1.925 | 6.0E-05 |
| *CBLN1* | -1.920 | 3.1E-24 |
| *CCL2* | -1.919 | 6.3E-05 |

| | | |
|---|---|---|
| *KCNA6* | -1.915 | 3.5E-10 |
| *C5orf49* | -1.912 | 1.4E-09 |
| *IRX1* | -1.898 | 1.8E-04 |
| *EFCC1* | -1.898 | 8.7E-04 |
| *TMEM179* | -1.884 | 8.7E-07 |
| *GALNT17* | -1.883 | 1.2E-04 |
| *GAD1* | -1.881 | 2.9E-03 |
| *THY1* | -1.876 | 3.9E-07 |
| *PCDH15* | -1.875 | 4.2E-06 |
| *PREX1* | -1.869 | 1.7E-04 |
| *ITGA11* | -1.861 | 3.8E-10 |
| *VWC2* | -1.860 | 1.9E-06 |
| *AMTN* | -1.856 | 7.3E-07 |
| *TEKT2* | -1.852 | 3.4E-10 |
| *CABP7* | -1.837 | 8.6E-06 |
| *LRRTM2* | -1.837 | 1.1E-03 |
| *CNIH2* | -1.828 | 2.5E-16 |
| *SLC8A3* | -1.828 | 3.5E-08 |
| *APC2* | -1.827 | 6.0E-07 |
| *IRX2* | -1.826 | 2.7E-04 |
| *IFI44L* | -1.824 | 2.6E-03 |
| *FAM107A* | -1.820 | 2.0E-02 |
| *RELN* | -1.818 | 6.5E-13 |
| *ZIC4* | -1.813 | 1.5E-03 |
| *LRP8* | -1.810 | 6.8E-11 |
| *FOSB* | -1.800 | 1.3E-05 |
| *GRM4* | -1.796 | 5.8E-09 |
| *TMEM178B* | -1.795 | 1.9E-13 |
| *KAAG1* | -1.793 | 3.1E-03 |
| *CFAP157* | -1.789 | 4.0E-04 |
| *PDZD4* | -1.786 | 2.5E-11 |
| *GRIA1* | -1.776 | 8.7E-08 |
| *SEPTIN3* | -1.775 | 1.4E-05 |
| *STK32B* | -1.773 | 9.0E-06 |
| *B4GALNT1* | -1.767 | 1.2E-05 |
| *LEFTY1* | -1.764 | 6.9E-10 |
| *FGF8* | -1.762 | 1.2E-03 |
| *ENO4* | -1.762 | 2.4E-05 |

| | | |
|---|---|---|
| *VAX1* | -1.756 | 1.0E-02 |
| *CD44* | -1.754 | 6.6E-16 |
| *RERG* | -1.746 | 1.1E-10 |
| *NUPR1* | -1.742 | 1.6E-02 |
| *SLC32A1* | -1.741 | 1.1E-03 |
| *DCLK1* | -1.734 | 6.0E-08 |
| *MEP1B* | -1.733 | 2.9E-07 |
| *STMN4* | -1.729 | 2.9E-04 |
| *GPM6A* | -1.722 | 5.8E-03 |
| *EVA1C* | -1.721 | 1.9E-03 |
| *SOX1* | -1.720 | 2.4E-04 |
| *SLC6A15* | -1.719 | 5.4E-14 |
| *PNMA8C* | -1.717 | 4.7E-04 |
| *CRB2* | -1.710 | 6.5E-04 |
| *CSPG5* | -1.709 | 3.5E-05 |
| *MMP7* | -1.708 | 1.5E-02 |
| *OTX2* | -1.706 | 4.6E-03 |
| *PCSK1N* | -1.705 | 7.0E-06 |
| *CXCR4* | -1.702 | 1.8E-11 |
| *FGFBP3* | -1.698 | 2.6E-05 |
| *LGALS1* | -1.696 | 2.2E-08 |
| *PTGER3* | -1.695 | 3.2E-09 |
| *GFRA3* | -1.689 | 8.2E-15 |
| *ARC* | -1.684 | 2.9E-03 |
| *TMPRSS5* | -1.680 | 3.7E-05 |
| *FIBIN* | -1.676 | 2.9E-04 |
| *KCNN3* | -1.676 | 8.0E-05 |
| *CHST8* | -1.674 | 1.3E-06 |
| *ILDR2* | -1.674 | 5.8E-04 |
| *CCDC74B* | -1.673 | 1.6E-17 |
| *HAPLN1* | -1.672 | 1.3E-12 |
| *POSTN* | -1.667 | 3.4E-06 |
| *NCAN* | -1.667 | 1.8E-02 |
| *NFIX* | -1.667 | 1.8E-06 |
| *PCDHGB1* | -1.663 | 1.7E-03 |
| *MAP3K7CL* | -1.658 | 5.2E-07 |
| *SPEF1* | -1.658 | 3.6E-05 |
| *ANKRD63* | -1.649 | 8.4E-05 |

| | | |
|---|---|---|
| *SULF1* | -1.644 | 4.2E-05 |
| *FEZ1* | -1.641 | 5.1E-07 |
| *PDE4B* | -1.630 | 5.5E-05 |
| *ECEL1* | -1.624 | 2.4E-13 |
| *SOGA3* | -1.624 | 2.8E-05 |
| *PTX3* | -1.622 | 2.7E-05 |
| *ADGRL4* | -1.619 | 8.1E-06 |
| *RHOJ* | -1.605 | 1.6E-06 |
| *GABRG3* | -1.604 | 5.8E-05 |
| *KL* | -1.598 | 1.3E-05 |
| *PTCH2* | -1.598 | 2.8E-03 |
| *MAB21L2* | -1.591 | 1.6E-02 |
| *CLEC18B* | -1.590 | 2.8E-07 |
| *NPHS1* | -1.589 | 3.6E-09 |
| *TMEM132B* | -1.587 | 8.5E-11 |
| *ADAMTS3* | -1.585 | 2.2E-04 |
| *NFIA* | -1.583 | 7.0E-07 |
| *CALN1* | -1.581 | 1.6E-04 |
| *SPON1* | -1.575 | 1.2E-02 |
| *HGFAC* | -1.566 | 6.9E-07 |
| *SCG5* | -1.564 | 8.6E-07 |
| *DLX5* | -1.561 | 1.7E-03 |
| *LRRIQ1* | -1.559 | 2.6E-03 |
| *NOVA2* | -1.558 | 1.3E-04 |
| *DMBX1* | -1.556 | 6.4E-06 |
| *PLPPR4* | -1.553 | 1.3E-03 |
| *PLA2G3* | -1.551 | 8.1E-04 |
| *VGF* | -1.547 | 6.6E-05 |
| *SYT5* | -1.544 | 1.6E-08 |
| *DGKG* | -1.542 | 4.7E-02 |
| *RSPO3* | -1.541 | 7.9E-03 |
| *LMO2* | -1.534 | 9.5E-04 |
| *VEGFC* | -1.533 | 4.9E-04 |
| *DCLK2* | -1.532 | 1.4E-06 |
| *CCDC184* | -1.532 | 3.3E-04 |
| *GNG2* | -1.531 | 4.1E-08 |
| *CSMD2* | -1.530 | 1.4E-05 |
| *CCDC81* | -1.529 | 1.7E-06 |

| | | |
|---|---|---|
| *PPM1E* | -1.524 | 4.6E-05 |
| *PLXNB3* | -1.520 | 1.4E-03 |
| *JAM2* | -1.519 | 3.0E-05 |
| *PTPRN* | -1.518 | 1.6E-16 |
| *NPAS3* | -1.513 | 2.0E-04 |
| *TRPC4* | -1.508 | 1.5E-04 |
| *ARMC3* | -1.508 | 1.1E-05 |
| *ATP8A2* | -1.507 | 8.5E-05 |
| *COL6A3* | -1.506 | 9.6E-06 |
| *CSPG4* | -1.504 | 1.7E-07 |
| *DCDC1* | -1.504 | 4.9E-06 |
| *PLCXD3* | -1.504 | 2.5E-11 |
| *TNC* | -1.497 | 6.8E-06 |
| *DNAAF3* | -1.493 | 1.5E-04 |
| *MIAT* | -1.485 | 2.6E-10 |
| *SHANK1* | -1.484 | 1.2E-02 |
| *ADAMTS14* | -1.484 | 3.2E-06 |
| *ADGRF5* | -1.483 | 3.7E-12 |
| *PRODH2* | -1.482 | 2.3E-04 |
| *SIX3* | -1.480 | 1.6E-03 |
| *PGM5* | -1.480 | 2.3E-07 |
| *OLFM1* | -1.478 | 2.8E-03 |
| *WFDC1* | -1.478 | 1.2E-02 |
| *DNAH6* | -1.478 | 2.8E-04 |
| *DKK2* | -1.476 | 2.0E-04 |
| *GRIN2B* | -1.472 | 3.3E-05 |
| *CADM3* | -1.467 | 4.8E-05 |
| *DNAAF1* | -1.467 | 1.4E-03 |
| *SHISAL1* | -1.466 | 3.0E-02 |
| *FGL1* | -1.463 | 2.0E-03 |
| *LRRC4* | -1.457 | 1.3E-06 |
| *MLIP* | -1.454 | 1.6E-03 |
| *GREM1* | -1.452 | 2.0E-12 |
| *DPYSL5* | -1.452 | 4.8E-04 |
| *CCDC74A* | -1.452 | 2.1E-05 |
| *TTYH1* | -1.451 | 1.6E-06 |
| *AK5* | -1.449 | 1.4E-04 |
| *CELF5* | -1.446 | 3.5E-03 |

| | | |
|---|---|---|
| *ALKAL2* | -1.445 | 5.4E-03 |
| *THBS2* | -1.444 | 2.6E-04 |
| *NOG* | -1.443 | 1.4E-03 |
| *NTRK3* | -1.440 | 1.8E-02 |
| *NLGN4X* | -1.437 | 3.5E-08 |
| *DHRS2* | -1.436 | 2.8E-18 |
| *SLIT1* | -1.435 | 4.4E-02 |
| *DLK1* | -1.434 | 3.9E-18 |
| *HMGCLL1* | -1.431 | 3.3E-08 |
| *KCNMB1* | -1.431 | 2.8E-03 |
| *FOXD1* | -1.428 | 3.4E-02 |
| *NKX2-1* | -1.426 | 2.3E-02 |
| *METRN* | -1.425 | 1.1E-15 |
| *ATP1A2* | -1.422 | 2.3E-03 |
| *IGSF9B* | -1.417 | 8.8E-07 |
| *DLX2* | -1.415 | 1.1E-02 |
| *RN7SL2* | -1.414 | 3.5E-04 |
| *NPTX1* | -1.414 | 1.3E-02 |
| *NCALD* | -1.410 | 7.8E-05 |
| *CYP26A1* | -1.402 | 3.4E-04 |
| *NCAM1* | -1.402 | 5.1E-07 |
| *CLEC11A* | -1.402 | 2.8E-04 |
| *ZEB1* | -1.401 | 5.6E-03 |
| *TMEM130* | -1.394 | 5.5E-03 |
| *CNTNAP3* | -1.392 | 7.4E-05 |
| *CHST7* | -1.390 | 4.6E-05 |
| *DYRK3* | -1.389 | 3.4E-07 |
| *RNF182* | -1.385 | 4.8E-05 |
| *CCK* | -1.384 | 1.3E-04 |
| *RGMA* | -1.380 | 2.2E-03 |
| *DLL3* | -1.379 | 3.2E-02 |
| *ZNF521* | -1.378 | 5.7E-04 |
| *TMSB15A* | -1.376 | 2.0E-03 |
| *ADAMTSL1* | -1.375 | 1.2E-03 |
| *GFAP* | -1.373 | 4.3E-04 |
| *DNER* | -1.373 | 8.4E-05 |
| *NIM1K* | -1.371 | 1.1E-03 |
| *TACR1* | -1.371 | 8.1E-06 |

| | | |
|---|---|---|
| *BEX1* | -1.367 | 2.9E-18 |
| *TMEM145* | -1.364 | 1.7E-05 |
| *CDH10* | -1.361 | 4.8E-12 |
| *CDO1* | -1.360 | 4.8E-05 |
| *LDB2* | -1.358 | 3.3E-02 |
| *ATOH8* | -1.356 | 9.0E-04 |
| *COL9A1* | -1.352 | 3.8E-03 |
| *LRRC10B* | -1.349 | 6.4E-05 |
| *SCN7A* | -1.347 | 1.2E-03 |
| *ST6GAL2* | -1.343 | 5.6E-04 |
| *PLPPR5* | -1.343 | 2.0E-05 |
| *SEMA5B* | -1.342 | 8.8E-04 |
| *KCNQ2* | -1.332 | 8.2E-03 |
| *BBOX1* | -1.332 | 9.2E-04 |
| *SPOCK3* | -1.329 | 2.5E-05 |
| *MAPK10* | -1.329 | 2.2E-05 |
| *NECAB2* | -1.328 | 4.7E-02 |
| *BCL2* | -1.324 | 4.4E-03 |
| *ACTG2* | -1.324 | 5.6E-05 |
| *PRR7* | -1.324 | 2.8E-07 |
| *SHC4* | -1.320 | 4.1E-06 |
| *FBXL16* | -1.315 | 5.4E-06 |
| *OLFML2B* | -1.310 | 1.2E-02 |
| *MAP2* | -1.309 | 1.6E-13 |
| *CCN3* | -1.307 | 1.1E-07 |
| *CTXN1* | -1.306 | 1.7E-10 |
| *PRRX1* | -1.305 | 9.5E-03 |
| *CALY* | -1.304 | 2.1E-05 |
| *ADGRV1* | -1.303 | 2.0E-08 |
| *RAPGEF4* | -1.300 | 7.5E-10 |
| *ZNF506* | -1.299 | 2.4E-08 |
| *ADAM12* | -1.296 | 3.8E-03 |
| *EPHB1* | -1.296 | 2.6E-04 |
| *HOPX* | -1.294 | 4.3E-04 |
| *RPRM* | -1.293 | 2.8E-04 |
| *NNAT* | -1.290 | 4.3E-09 |
| *CACNG8* | -1.288 | 1.4E-04 |
| *DIRAS2* | -1.286 | 3.1E-03 |

| | | |
|---|---|---|
| *LOXL2* | -1.286 | 4.4E-11 |
| *KISS1R* | -1.282 | 5.4E-06 |
| *FRZB* | -1.280 | 1.1E-02 |
| *HS6ST3* | -1.278 | 7.2E-13 |
| *LRFN2* | -1.277 | 4.6E-03 |
| *DOK5* | -1.276 | 3.9E-02 |
| *PACRG* | -1.273 | 5.6E-07 |
| *SOX2* | -1.271 | 4.0E-03 |
| *NRXN2* | -1.269 | 2.4E-02 |
| *ISLR2* | -1.267 | 3.2E-02 |
| *SCUBE2* | -1.265 | 1.3E-02 |
| *SYT11* | -1.262 | 3.2E-07 |
| *LRRC4B* | -1.261 | 1.0E-02 |
| *UNC5A* | -1.260 | 7.3E-03 |
| *DENND2A* | -1.258 | 1.9E-04 |
| *FGFR1* | -1.258 | 9.3E-13 |
| *GPC2* | -1.257 | 4.6E-11 |
| *PITPNC1* | -1.253 | 1.6E-12 |
| *NUF2* | -1.251 | 1.4E-04 |
| *COL8A1* | -1.248 | 4.5E-03 |
| *ASTN1* | -1.247 | 5.1E-03 |
| *TOX2* | -1.241 | 3.5E-02 |
| *PNMA2* | -1.240 | 3.1E-26 |
| *DNAH7* | -1.238 | 1.8E-03 |
| *STARD9* | -1.238 | 1.3E-04 |
| *RTN1* | -1.235 | 1.3E-04 |
| *MAP6* | -1.235 | 3.7E-07 |
| *RGS16* | -1.226 | 9.1E-08 |
| *FAM124A* | -1.226 | 3.4E-02 |
| *PNMT* | -1.223 | 9.8E-05 |
| *ADCYAP1R1* | -1.222 | 1.6E-03 |
| *BGN* | -1.222 | 3.1E-03 |
| *RHPN1* | -1.218 | 1.5E-06 |
| *FAM72B* | -1.217 | 5.7E-03 |
| *VIM* | -1.216 | 5.2E-07 |
| *PIF1* | -1.213 | 8.7E-05 |
| *PAK3* | -1.212 | 1.8E-07 |
| *SALL1* | -1.212 | 2.8E-02 |

| | | |
|---|---|---|
| *UBXN10* | -1.210 | 4.1E-07 |
| *BAIAP3* | -1.210 | 6.1E-14 |
| *CCDC160* | -1.207 | 1.5E-03 |
| *KCNJ11* | -1.205 | 8.7E-15 |
| *ELAVL4* | -1.198 | 3.2E-06 |
| *CFTR* | -1.197 | 5.0E-04 |
| *PCDHGC4* | -1.197 | 9.8E-03 |
| *FOS* | -1.197 | 3.8E-05 |
| *CCDC181* | -1.194 | 2.4E-04 |
| *ELAVL3* | -1.191 | 5.2E-03 |
| *NLGN3* | -1.191 | 2.7E-04 |
| *SYNPR* | -1.191 | 5.6E-03 |
| *AMHR2* | -1.190 | 7.2E-04 |
| *NKD1* | -1.190 | 2.6E-02 |
| *GDAP1L1* | -1.189 | 2.4E-04 |
| *CHRDL1* | -1.187 | 3.7E-02 |
| *SUGCT* | -1.186 | 1.1E-03 |
| *MATN2* | -1.184 | 5.2E-07 |
| *SLC35F1* | -1.184 | 4.9E-03 |
| *COL9A3* | -1.183 | 1.0E-06 |
| *SFTA1P* | -1.181 | 9.6E-03 |
| *MDH1B* | -1.180 | 1.6E-04 |
| *FAM181B* | -1.178 | 2.4E-04 |
| *GREM2* | -1.176 | 3.6E-02 |
| *FXYD2* | -1.175 | 4.6E-09 |
| *GHRL* | -1.174 | 7.9E-04 |
| *CHRNA5* | -1.173 | 6.0E-07 |
| *S1PR3* | -1.172 | 2.0E-05 |
| *HIC1* | -1.172 | 2.0E-04 |
| *ANGPTL4* | -1.171 | 1.1E-05 |
| *PAPPA* | -1.171 | 7.2E-05 |
| *KIF5A* | -1.171 | 1.9E-05 |
| *SUSD4* | -1.170 | 6.5E-05 |
| *KRT4* | -1.170 | 2.7E-03 |
| *SYT2* | -1.170 | 2.3E-05 |
| *LMO3* | -1.168 | 1.4E-03 |
| *CRMP1* | -1.168 | 3.0E-06 |
| *JAM3* | -1.166 | 5.7E-04 |

| | | |
|---|---|---|
| *SCG3* | -1.164 | 9.4E-23 |
| *OIP5* | -1.164 | 2.6E-03 |
| *SOX21* | -1.163 | 2.5E-02 |
| *HSF4* | -1.162 | 1.6E-06 |
| *TAGLN3* | -1.158 | 6.7E-03 |
| *DCHS2* | -1.157 | 1.2E-02 |
| *LDLRAD4* | -1.156 | 9.4E-09 |
| *RNF165* | -1.156 | 2.1E-02 |
| *CORO1A* | -1.154 | 4.6E-06 |
| *FAM131C* | -1.154 | 1.6E-06 |
| *ANXA10* | -1.154 | 1.5E-02 |
| *FHL1* | -1.153 | 5.1E-04 |
| *FAM201A* | -1.151 | 2.1E-02 |
| *SH2D3C* | -1.151 | 2.4E-02 |
| *GLIPR1* | -1.150 | 7.1E-05 |
| *FBLL1* | -1.146 | 3.4E-05 |
| *IGSF11* | -1.141 | 3.4E-05 |
| *CHN1* | -1.140 | 1.1E-04 |
| *CENPW* | -1.140 | 4.3E-07 |
| *MAPK11* | -1.139 | 2.7E-04 |
| *SLC6A2* | -1.138 | 5.1E-03 |
| *NR2F1* | -1.136 | 3.5E-03 |
| *NALCN* | -1.135 | 1.0E-03 |
| *TSPOAP1* | -1.134 | 5.6E-09 |
| *NRXN1* | -1.134 | 8.8E-09 |
| *LRRC24* | -1.132 | 6.0E-05 |
| *NAP1L3* | -1.132 | 2.6E-06 |
| *DRC7* | -1.131 | 2.5E-04 |
| *WNT10A* | -1.131 | 3.3E-04 |
| *SYNDIG1* | -1.130 | 3.3E-02 |
| *STMN2* | -1.128 | 3.8E-19 |
| *PEG13* | -1.124 | 5.1E-04 |
| *MEX3B* | -1.124 | 4.8E-04 |
| *EGR1* | -1.123 | 1.7E-03 |
| *TRIM9* | -1.123 | 3.3E-03 |
| *UNC13A* | -1.122 | 6.2E-17 |
| *KCNMA1* | -1.121 | 2.4E-09 |
| *CYP1B1* | -1.118 | 3.8E-05 |

| | | |
|---|---|---|
| *PPIL6* | -1.117 | 1.1E-04 |
| *RRAD* | -1.117 | 7.4E-04 |
| *DPYSL4* | -1.117 | 8.7E-08 |
| *ABCB4* | -1.116 | 1.6E-02 |
| *BMERB1* | -1.115 | 2.4E-07 |
| *RAB36* | -1.114 | 3.7E-08 |
| *ADGRB3* | -1.109 | 2.6E-05 |
| *ST8SIA2* | -1.105 | 1.1E-03 |
| *CCNA1* | -1.105 | 5.8E-05 |
| *GFPT2* | -1.104 | 4.2E-03 |
| *GRIN2A* | -1.101 | 1.4E-05 |
| *EFNB3* | -1.101 | 8.3E-06 |
| *ETV1* | -1.097 | 1.8E-04 |
| *GRM8* | -1.096 | 9.9E-04 |
| *TMEM59L* | -1.094 | 1.4E-04 |
| *RGS20* | -1.093 | 1.6E-02 |
| *C8A* | -1.093 | 2.4E-03 |
| *ARHGAP33* | -1.091 | 2.9E-04 |
| *ZEB2* | -1.091 | 4.9E-02 |
| *RAB33A* | -1.089 | 3.1E-02 |
| *LHFPL6* | -1.088 | 1.9E-06 |
| *TMEFF1* | -1.088 | 5.0E-03 |
| *ELAVL2* | -1.088 | 4.6E-03 |
| *HCN1* | -1.088 | 1.1E-05 |
| *DPF1* | -1.087 | 2.4E-03 |
| *FOSL1* | -1.083 | 1.3E-02 |
| *ATCAY* | -1.083 | 3.5E-03 |
| *EDN3* | -1.083 | 1.0E-03 |
| *PNMA8B* | -1.081 | 5.4E-06 |
| *KCNIP1* | -1.081 | 4.4E-03 |
| *ACTA2* | -1.081 | 1.0E-03 |
| *NEK10* | -1.079 | 4.2E-02 |
| *MAPK8IP1* | -1.079 | 2.5E-13 |
| *SYDE1* | -1.079 | 1.2E-06 |
| *NTM* | -1.077 | 5.1E-05 |
| *TMEM262* | -1.076 | 1.8E-04 |
| *KRT20* | -1.075 | 2.4E-02 |
| *DRP2* | -1.074 | 1.2E-02 |

| | | |
|---|---|---|
| *HOXB2* | -1.073 | 4.3E-02 |
| *CYTL1* | -1.071 | 1.8E-03 |
| *HAGHL* | -1.071 | 1.0E-04 |
| *DIRAS3* | -1.071 | 4.3E-05 |
| *PLCH2* | -1.070 | 3.4E-04 |
| *DGKI* | -1.068 | 1.7E-02 |
| *LHFPL4* | -1.068 | 6.0E-05 |
| *ZCCHC18* | -1.065 | 8.1E-07 |
| *SOSTDC1* | -1.061 | 2.9E-02 |
| *CADPS* | -1.061 | 7.4E-06 |
| *SLC15A3* | -1.061 | 4.9E-02 |
| *FAT3* | -1.058 | 1.0E-03 |
| *NANOS1* | -1.058 | 8.3E-05 |
| *KIAA1549L* | -1.057 | 4.0E-02 |
| *CCDC78* | -1.054 | 3.0E-03 |
| *HS3ST3B1* | -1.054 | 1.6E-05 |
| *INA* | -1.053 | 6.2E-12 |
| *FZD9* | -1.052 | 2.4E-02 |
| *RSPH4A* | -1.051 | 2.4E-05 |
| *TMEM151B* | -1.051 | 6.7E-03 |
| *LGR5* | -1.051 | 6.0E-04 |
| *GPR68* | -1.048 | 8.6E-04 |
| *PHF21B* | -1.048 | 3.4E-02 |
| *DRAXIN* | -1.046 | 4.1E-02 |
| *SIX2* | -1.045 | 1.2E-05 |
| *TSNAXIP1* | -1.044 | 3.5E-04 |
| *CENPF* | -1.044 | 2.5E-05 |
| *ANO4* | -1.043 | 5.5E-11 |
| *SYT4* | -1.042 | 5.2E-07 |
| *SCRG1* | -1.038 | 1.3E-02 |
| *PCDHB15* | -1.038 | 1.0E-02 |
| *EFCAB12* | -1.038 | 6.8E-03 |
| *TMEM121* | -1.037 | 8.6E-03 |
| *TENM2* | -1.037 | 1.3E-05 |
| *CRYAB* | -1.035 | 5.0E-03 |
| *TRIM46* | -1.034 | 6.1E-05 |
| *PSRC1* | -1.030 | 5.2E-05 |
| *HSPA6* | -1.027 | 3.1E-02 |

| | | |
|---|---|---|
| *ZNF350* | -1.026 | 5.2E-04 |
| *ATP1A3* | -1.025 | 4.8E-02 |
| *XKR4* | -1.025 | 9.7E-05 |
| *EDAR* | -1.025 | 2.4E-03 |
| *ARHGEF6* | -1.022 | 2.1E-02 |
| *MAPK4* | -1.021 | 5.7E-10 |
| *MRC2* | -1.021 | 6.7E-08 |
| *SOD3* | -1.020 | 5.1E-05 |
| *PCYT1B* | -1.020 | 4.3E-06 |
| *REC8* | -1.020 | 5.3E-20 |
| *FAM167A* | -1.019 | 3.1E-09 |
| *WDR25* | -1.019 | 3.5E-12 |
| *CACNG7* | -1.018 | 3.9E-02 |
| *TNR* | -1.017 | 2.0E-09 |
| *UNC5D* | -1.016 | 3.0E-05 |
| *PHACTR1* | -1.016 | 1.2E-02 |
| *CCDC9B* | -1.015 | 3.1E-02 |
| *LGI3* | -1.015 | 4.1E-02 |
| *INSYN1* | -1.012 | 7.5E-03 |
| *RNF150* | -1.012 | 1.4E-04 |
| *FJX1* | -1.011 | 5.2E-11 |
| *GALNT15* | -1.009 | 1.3E-03 |
| *LRRC49* | -1.008 | 3.7E-05 |
| *WDR97* | -1.007 | 4.3E-05 |
| *STMN3* | -1.007 | 1.1E-04 |
| *NOTCH1* | -1.006 | 1.5E-06 |
| *PCDHB5* | -1.005 | 1.9E-03 |
| *TPM2* | -1.003 | 9.9E-07 |

**Table S7.** Top differentially expressed genes (DEGs) ($Log_2$ FC > 1, $p < 0.05$) upregulated in *RFX3* KO1 and *RFX3* KO2 at islet stage (S6) compared to WT.

| **Gene ID** | **$Log_2$ Fold Change** | ***p*-value** |
|---|---|---|
| *ADIRF* | 3.393 | 4.8E-02 |
| *MT1H* | 3.333 | 2.3E-05 |
| *ALOX15B* | 3.250 | 2.1E-03 |
| *CPN2* | 2.992 | 2.4E-03 |
| *ADH4* | 2.967 | 7.0E-16 |
| *ZPLD1* | 2.889 | 2.6E-15 |
| *C17orf99* | 2.805 | 1.1E-03 |
| *PDZK1IP1* | 2.801 | 1.0E-04 |
| *GABRG1* | 2.748 | 9.1E-09 |
| *XPNPEP2* | 2.740 | 3.7E-03 |
| *TBX4* | 2.724 | 5.8E-03 |
| *SLC14A2* | 2.598 | 3.4E-26 |
| *SUSD2* | 2.576 | 5.5E-03 |
| *SAA1* | 2.569 | 1.2E-03 |
| *SLC10A1* | 2.513 | 2.0E-03 |
| *AGXT2* | 2.482 | 3.6E-03 |
| *CPS1* | 2.439 | 2.9E-03 |
| *ODAPH* | 2.366 | 1.7E-03 |
| *ALDH1L1* | 2.359 | 1.2E-02 |
| *SOWAHD* | 2.325 | 5.4E-03 |
| *PCP4L1* | 2.287 | 2.2E-04 |
| *KNG1* | 2.260 | 7.1E-04 |
| *SULT2A1* | 2.247 | 1.1E-02 |
| *P2RY4* | 2.151 | 2.5E-03 |
| *CSF3R* | 2.141 | 9.6E-03 |
| *CHP2* | 2.112 | 8.6E-03 |
| *SLC6A20* | 2.104 | 5.1E-03 |
| *SOAT2* | 2.098 | 1.6E-02 |
| *SLC5A8* | 2.081 | 6.2E-05 |
| *MSLN* | 2.073 | 1.5E-02 |
| *C4BPA* | 2.072 | 4.4E-03 |
| *ORM1* | 2.069 | 3.1E-03 |
| *TNFRSF1B* | 2.055 | 6.5E-03 |
| *MAT1A* | 2.040 | 5.6E-04 |
| *PRSS36* | 2.034 | 1.3E-04 |

| | | |
|---|---|---|
| *ABCC2* | 2.029 | 8.9E-03 |
| *ORM2* | 2.028 | 2.2E-03 |
| *KCNE3* | 2.024 | 6.4E-05 |
| *THPO* | 2.019 | 6.3E-04 |
| *SLC38A5* | 2.010 | 1.9E-03 |
| *ABI3BP* | 1.999 | 4.7E-03 |
| *APELA* | 1.993 | 6.9E-03 |
| *NPL* | 1.970 | 9.2E-04 |
| *CASP5* | 1.963 | 8.9E-03 |
| *XKRX* | 1.960 | 3.4E-03 |
| *CD244* | 1.956 | 2.9E-03 |
| *S100G* | 1.939 | 1.7E-02 |
| *CTSV* | 1.928 | 8.2E-03 |
| *OTOP2* | 1.919 | 2.5E-02 |
| *ATP6V0A4* | 1.914 | 6.3E-03 |
| *SLC46A3* | 1.913 | 3.7E-03 |
| *MS4A10* | 1.908 | 4.5E-02 |
| *DKK4* | 1.908 | 3.6E-03 |
| *ERVH48-1* | 1.904 | 9.5E-05 |
| *AIRE* | 1.903 | 2.6E-03 |
| *BST2* | 1.901 | 4.8E-03 |
| *ABCA10* | 1.871 | 5.0E-04 |
| *DEFB1* | 1.869 | 4.9E-03 |
| *APOB* | 1.867 | 3.9E-03 |
| *TPH1* | 1.863 | 1.9E-17 |
| *IL22RA1* | 1.862 | 8.8E-03 |
| *LYPD2* | 1.850 | 7.5E-06 |
| *P2RX1* | 1.848 | 3.4E-03 |
| *SLPI* | 1.848 | 3.0E-03 |
| *MUC17* | 1.847 | 3.4E-03 |
| *SLC23A1* | 1.833 | 6.2E-03 |
| *APOD* | 1.823 | 2.7E-03 |
| *PAPPA2* | 1.822 | 8.3E-14 |
| *SAA2* | 1.818 | 8.6E-03 |
| *CTXND1* | 1.816 | 8.2E-04 |
| *AOC1* | 1.816 | 9.9E-05 |
| *G0S2* | 1.815 | 1.1E-02 |
| *ALX1* | 1.809 | 1.2E-03 |

| | | |
|---|---|---|
| *KCNA4* | 1.808 | 2.4E-06 |
| *GBA3* | 1.805 | 1.2E-02 |
| *TRIM31* | 1.805 | 1.3E-02 |
| *TF* | 1.798 | 3.5E-02 |
| *MRO* | 1.795 | 5.7E-03 |
| *PSAPL1* | 1.795 | 4.5E-04 |
| *CPA2* | 1.793 | 8.8E-03 |
| *KCNJ13* | 1.791 | 1.8E-04 |
| *CASP1* | 1.787 | 2.0E-03 |
| *MLN* | 1.780 | 1.8E-02 |
| *INMT* | 1.777 | 4.1E-02 |
| *ENPP7* | 1.766 | 4.8E-02 |
| *CCR1* | 1.755 | 7.2E-04 |
| *ENPP3* | 1.749 | 1.5E-02 |
| *KCNJ12* | 1.735 | 4.5E-03 |
| *CLCA1* | 1.725 | 3.7E-02 |
| *KLK1* | 1.716 | 1.6E-02 |
| *LRRC19* | 1.697 | 5.3E-03 |
| *CKM* | 1.694 | 2.8E-08 |
| *FADS6* | 1.692 | 2.1E-02 |
| *ERICH4* | 1.690 | 2.0E-02 |
| *TMEM171* | 1.687 | 1.3E-02 |
| *MOGAT2* | 1.686 | 3.5E-02 |
| *WIPF3* | 1.685 | 1.3E-04 |
| *OTOP3* | 1.684 | 1.8E-02 |
| *SLC30A2* | 1.676 | 1.9E-02 |
| *TDO2* | 1.663 | 3.1E-02 |
| *CREG2* | 1.652 | 1.0E-02 |
| *RBP4* | 1.645 | 2.9E-02 |
| *SLC15A1* | 1.641 | 6.8E-03 |
| *UGT2B17* | 1.641 | 1.7E-02 |
| *APOL3* | 1.640 | 3.1E-02 |
| *CSF2RA* | 1.639 | 3.0E-08 |
| *ITIH3* | 1.639 | 2.2E-02 |
| *PRKG2* | 1.635 | 1.9E-02 |
| *EGF* | 1.630 | 2.0E-02 |
| *COL14A1* | 1.628 | 1.3E-02 |
| *PIGR* | 1.627 | 3.4E-03 |

| | | |
|---|---|---|
| *TXNIP* | 1.620 | 5.0E-05 |
| *AKR1D1* | 1.617 | 7.1E-06 |
| *FOLH1* | 1.612 | 1.4E-02 |
| *ANPEP* | 1.611 | 8.7E-03 |
| *CD3G* | 1.610 | 8.1E-03 |
| *MUCL3* | 1.610 | 9.6E-09 |
| *HMCN2* | 1.608 | 1.6E-02 |
| *GPA33* | 1.603 | 4.6E-03 |
| *IL3RA* | 1.601 | 2.0E-02 |
| *NOX1* | 1.600 | 1.6E-03 |
| *TFEC* | 1.580 | 1.2E-02 |
| *ZG16* | 1.579 | 1.0E-02 |
| *RGPD1* | 1.574 | 8.0E-04 |
| *CD68* | 1.573 | 1.2E-02 |
| *FGF23* | 1.573 | 4.1E-02 |
| *LY75* | 1.571 | 2.6E-04 |
| *RUBCNL* | 1.571 | 6.7E-03 |
| *SLC26A3* | 1.562 | 2.9E-02 |
| *OLR1* | 1.557 | 4.1E-04 |
| *NAGS* | 1.554 | 5.9E-03 |
| *C3* | 1.545 | 7.1E-09 |
| *MBNL3* | 1.545 | 1.4E-03 |
| *FAM3D* | 1.545 | 5.0E-03 |
| *SHBG* | 1.535 | 4.6E-03 |
| *FCGBP* | 1.534 | 2.7E-02 |
| *PPP1R3G* | 1.533 | 2.1E-03 |
| *SMLR1* | 1.529 | 1.5E-02 |
| *TRIM63* | 1.519 | 1.7E-02 |
| *DGAT2* | 1.518 | 2.8E-02 |
| *STARD5* | 1.507 | 2.0E-02 |
| *C2* | 1.506 | 2.1E-02 |
| *MOGAT3* | 1.497 | 1.8E-02 |
| *SLC6A19* | 1.495 | 2.9E-02 |
| *RHBG* | 1.492 | 3.0E-02 |
| *CLDN19* | 1.484 | 6.0E-03 |
| *SLC3A1* | 1.480 | 1.6E-02 |
| *ART5* | 1.478 | 2.2E-02 |
| *P2RY6* | 1.470 | 7.8E-03 |

| | | |
|---|---|---|
| *PRDM1* | 1.470 | 6.0E-04 |
| *FRMD1* | 1.469 | 1.3E-02 |
| *CCRL2* | 1.467 | 4.4E-04 |
| *KEL* | 1.466 | 1.2E-02 |
| *FAM151A* | 1.460 | 1.6E-02 |
| *MUC3A* | 1.457 | 4.3E-03 |
| *MELTF* | 1.452 | 2.0E-02 |
| *MUC2* | 1.441 | 1.4E-02 |
| *PAH* | 1.438 | 5.0E-07 |
| *TAC3* | 1.437 | 1.3E-02 |
| *BTNL8* | 1.434 | 4.7E-02 |
| *NR1I3* | 1.430 | 4.6E-02 |
| *NOS3* | 1.429 | 3.2E-02 |
| *CEACAM7* | 1.428 | 4.7E-04 |
| *SLC7A9* | 1.426 | 1.3E-02 |
| *CPXM2* | 1.425 | 2.4E-02 |
| *PIEZO2* | 1.424 | 1.7E-02 |
| *ASS1* | 1.412 | 1.8E-04 |
| *LGALS2* | 1.409 | 4.4E-02 |
| *ACE2* | 1.406 | 5.3E-04 |
| *GBP2* | 1.405 | 6.6E-09 |
| *SLC51B* | 1.399 | 6.2E-03 |
| *HYAL1* | 1.399 | 3.7E-03 |
| *DHDH* | 1.389 | 3.0E-02 |
| *SLC27A2* | 1.384 | 4.0E-04 |
| *TYRP1* | 1.384 | 1.4E-02 |
| *ZNF385B* | 1.382 | 1.3E-02 |
| *ADH1A* | 1.382 | 3.0E-11 |
| *MYL3* | 1.381 | 1.5E-02 |
| *RETREG1* | 1.379 | 3.8E-04 |
| *MYEOV* | 1.375 | 6.4E-04 |
| *CTSA* | 1.372 | 8.9E-03 |
| *BATF2* | 1.371 | 6.8E-03 |
| *TNFSF10* | 1.370 | 2.7E-03 |
| *MFSD2A* | 1.368 | 4.7E-03 |
| *APOA2* | 1.363 | 1.5E-03 |
| *GSTA2* | 1.359 | 3.7E-02 |
| *KCNJ5* | 1.356 | 8.9E-05 |

| | | |
|---|---|---|
| *BMP8B* | 1.354 | 3.7E-02 |
| *GPC3* | 1.352 | 4.3E-02 |
| *MUC13* | 1.351 | 1.1E-02 |
| *ROS1* | 1.347 | 3.7E-05 |
| *PBLD* | 1.345 | 1.5E-03 |
| *SP8* | 1.340 | 3.6E-02 |
| *C1orf115* | 1.337 | 5.4E-04 |
| *GPAT3* | 1.337 | 3.4E-04 |
| *RARRES1* | 1.332 | 4.1E-02 |
| *CDHR2* | 1.331 | 4.6E-02 |
| *MST1* | 1.329 | 3.2E-02 |
| *HSD3B1* | 1.327 | 8.5E-04 |
| *PI3* | 1.326 | 1.1E-02 |
| *SLC16A10* | 1.326 | 6.9E-04 |
| *MRGPRF* | 1.325 | 5.4E-03 |
| *ACTN3* | 1.325 | 2.8E-03 |
| *TMEM253* | 1.324 | 1.8E-02 |
| *KYNU* | 1.322 | 1.7E-05 |
| *UPB1* | 1.315 | 4.0E-02 |
| *ASAH2* | 1.310 | 3.0E-02 |
| *SLC23A3* | 1.307 | 1.8E-04 |
| *DUSP9* | 1.305 | 2.1E-03 |
| *CHST13* | 1.302 | 1.0E-02 |
| *SLC2A2* | 1.292 | 1.0E-03 |
| *TRIM10* | 1.290 | 3.9E-02 |
| *ABCG2* | 1.287 | 8.9E-04 |
| *PTCSC2* | 1.287 | 3.6E-02 |
| *APOBEC3D* | 1.287 | 1.2E-02 |
| *SLC18A2* | 1.286 | 4.9E-07 |
| *RAB42* | 1.285 | 5.5E-04 |
| *CEL* | 1.283 | 6.3E-03 |
| *AADAC* | 1.282 | 3.6E-02 |
| *CD302* | 1.273 | 4.0E-04 |
| *RAB29* | 1.273 | 1.1E-02 |
| *NCR3LG1* | 1.271 | 1.4E-03 |
| *SLC7A8* | 1.270 | 1.6E-03 |
| *STRIP2* | 1.267 | 6.6E-03 |
| *CDHR5* | 1.264 | 1.6E-02 |

| | | |
|---|---|---|
| *COMP* | 1.263 | 1.0E-07 |
| *KBTBD12* | 1.262 | 5.3E-03 |
| *FAM20A* | 1.261 | 1.4E-03 |
| *TIFA* | 1.259 | 9.1E-04 |
| *C10orf95* | 1.256 | 7.7E-03 |
| *TRPA1* | 1.255 | 2.9E-04 |
| *SULT1B1* | 1.254 | 2.0E-03 |
| *PIWIL2* | 1.254 | 2.2E-02 |
| *AFP* | 1.251 | 2.7E-02 |
| *PRSS3* | 1.250 | 3.5E-02 |
| *INSIG1* | 1.247 | 1.9E-02 |
| *CADM2* | 1.247 | 1.1E-10 |
| *SLCO4C1* | 1.242 | 4.7E-04 |
| *SLC35G1* | 1.240 | 1.4E-03 |
| *MALRD1* | 1.239 | 2.1E-02 |
| *APOBR* | 1.238 | 3.4E-02 |
| *CYP8B1* | 1.237 | 1.9E-04 |
| *TGM2* | 1.236 | 1.3E-02 |
| *SLC2A9* | 1.233 | 2.1E-02 |
| *FMOD* | 1.232 | 2.1E-02 |
| *TNFSF14* | 1.230 | 3.4E-02 |
| *MYO1A* | 1.230 | 3.6E-03 |
| *GGT5* | 1.228 | 3.9E-03 |
| *PLA1A* | 1.226 | 2.1E-02 |
| *TUBAL3* | 1.225 | 4.6E-02 |
| *SULT1C2* | 1.224 | 3.6E-04 |
| *MOCOS* | 1.221 | 7.5E-03 |
| *CYP2W1* | 1.221 | 1.5E-03 |
| *ADAMTSL4* | 1.215 | 1.8E-03 |
| *FUT2* | 1.213 | 2.5E-04 |
| *ITPKA* | 1.213 | 8.2E-03 |
| *GIPC2* | 1.211 | 1.9E-03 |
| *CYP51A1* | 1.208 | 1.2E-03 |
| *TMEFF2* | 1.204 | 1.8E-13 |
| *PKD1L2* | 1.202 | 9.3E-04 |
| *OAT* | 1.198 | 1.9E-02 |
| *CA2* | 1.198 | 1.7E-04 |
| *GSTA1* | 1.195 | 4.5E-02 |

| | | |
|---|---|---|
| *PRODH* | 1.195 | 3.9E-02 |
| *TPP1* | 1.194 | 9.6E-04 |
| *TMEM220* | 1.194 | 1.0E-02 |
| *LTF* | 1.192 | 1.3E-05 |
| *SLC31A2* | 1.192 | 6.3E-04 |
| *HHLA2* | 1.191 | 2.8E-02 |
| *FABP1* | 1.186 | 2.2E-03 |
| *MUC12* | 1.186 | 2.2E-04 |
| *ANKRD40CL* | 1.180 | 2.2E-02 |
| *ALPG* | 1.177 | 1.5E-02 |
| *OTC* | 1.173 | 3.9E-02 |
| *SPX* | 1.173 | 3.6E-03 |
| *XYLB* | 1.171 | 8.7E-03 |
| *ADGRF1* | 1.170 | 1.2E-05 |
| *ADGRG7* | 1.165 | 7.3E-03 |
| *MYBPHL* | 1.163 | 3.1E-02 |
| *TRIM55* | 1.159 | 2.2E-02 |
| *KCNS1* | 1.159 | 6.0E-04 |
| *PTK6* | 1.154 | 1.5E-02 |
| *EPHX4* | 1.152 | 2.5E-04 |
| *EBI3* | 1.152 | 5.1E-04 |
| *CD8B* | 1.149 | 1.8E-03 |
| *MSMO1* | 1.146 | 1.3E-03 |
| *METTL27* | 1.146 | 3.4E-02 |
| *CDH22* | 1.144 | 1.6E-03 |
| *TRPV3* | 1.141 | 2.7E-05 |
| *CYP27A1* | 1.141 | 5.3E-03 |
| *ALPK1* | 1.140 | 1.4E-03 |
| *GUCY2C* | 1.136 | 3.1E-03 |
| *SMPD3* | 1.135 | 4.8E-02 |
| *NPPB* | 1.133 | 4.9E-02 |
| *CIDEB* | 1.131 | 3.9E-02 |
| *CREB3L3* | 1.130 | 3.9E-03 |
| *PNPLA1* | 1.129 | 3.3E-02 |
| *CEBPA* | 1.126 | 3.6E-03 |
| *IHH* | 1.126 | 2.0E-03 |
| *PIP5K1B* | 1.125 | 9.7E-04 |
| *BNIP5* | 1.125 | 4.2E-03 |

| | | |
|---|---|---|
| *IRAG2* | 1.123 | 9.5E-03 |
| *TMEM52* | 1.123 | 2.3E-02 |
| *GK* | 1.114 | 5.1E-03 |
| *LGMN* | 1.111 | 3.3E-03 |
| *IL1R2* | 1.111 | 5.0E-03 |
| *MST1R* | 1.110 | 1.0E-03 |
| *CYP4V2* | 1.109 | 1.8E-04 |
| *F2* | 1.108 | 7.0E-03 |
| *LPGAT1* | 1.104 | 7.1E-04 |
| *KLHL13* | 1.104 | 9.9E-03 |
| *MAB21L3* | 1.103 | 9.5E-03 |
| *MGAM* | 1.099 | 1.4E-02 |
| *SEMA6D* | 1.099 | 1.2E-02 |
| *DHRS11* | 1.097 | 5.2E-03 |
| *STEAP2* | 1.090 | 1.2E-07 |
| *CRYBG1* | 1.088 | 3.1E-04 |
| *GATM* | 1.087 | 1.0E-02 |
| *FLVCR2* | 1.085 | 5.7E-03 |
| *JPH1* | 1.083 | 3.9E-03 |
| *TTC22* | 1.083 | 3.2E-03 |
| *MGST1* | 1.077 | 1.9E-03 |
| *CDC42EP2* | 1.076 | 1.5E-05 |
| *TMEM150B* | 1.074 | 3.6E-02 |
| *CDX1* | 1.073 | 8.3E-03 |
| *GPLD1* | 1.070 | 3.4E-04 |
| *PRSS1* | 1.069 | 7.8E-03 |
| *DAB1* | 1.069 | 1.5E-02 |
| *NR1H4* | 1.068 | 9.3E-03 |
| *LTK* | 1.067 | 2.5E-02 |
| *FOXE1* | 1.067 | 2.5E-02 |
| *GABRB1* | 1.064 | 2.9E-03 |
| *VDR* | 1.064 | 1.5E-03 |
| *DLX3* | 1.062 | 6.8E-03 |
| *NOD2* | 1.062 | 1.6E-02 |
| *IFIT3* | 1.061 | 5.5E-03 |
| *RAB3IL1* | 1.061 | 4.1E-02 |
| *ACE* | 1.060 | 7.4E-03 |
| *SMIM31* | 1.054 | 5.5E-03 |

| | | |
|---|---|---|
| *ALDOC* | 1.053 | 1.4E-02 |
| *FGA* | 1.053 | 8.6E-03 |
| *PCDHA10* | 1.049 | 2.5E-03 |
| *CASP10* | 1.047 | 3.1E-05 |
| *CLDN15* | 1.044 | 9.7E-04 |
| *TMEM92* | 1.043 | 1.6E-03 |
| *TMEM38B* | 1.041 | 6.0E-03 |
| *CARD14* | 1.041 | 7.4E-03 |
| *ALDH3B1* | 1.039 | 9.1E-03 |
| *SELENOP* | 1.039 | 1.2E-02 |
| *TRIM14* | 1.039 | 3.6E-03 |
| *TMEM144* | 1.036 | 9.5E-04 |
| *PDE11A* | 1.034 | 1.9E-02 |
| *B3GALT5* | 1.034 | 7.1E-04 |
| *CA4* | 1.033 | 3.5E-06 |
| *ALDH5A1* | 1.032 | 6.6E-04 |
| *VIPR1* | 1.032 | 2.8E-04 |
| *HSPE1-MOB4* | 1.027 | 2.5E-02 |
| *TFPI* | 1.027 | 1.3E-09 |
| *STOM* | 1.026 | 4.8E-04 |
| *MAF* | 1.026 | 3.8E-03 |
| *F7* | 1.026 | 1.8E-02 |
| *S100P* | 1.025 | 2.8E-04 |
| *BMP2K* | 1.025 | 1.3E-05 |
| *ERAP2* | 1.024 | 1.4E-04 |
| *STARD8* | 1.024 | 3.3E-02 |
| *OAF* | 1.023 | 1.3E-02 |
| *PRR15* | 1.021 | 7.6E-04 |
| *GALNT5* | 1.020 | 1.7E-04 |
| *BMP8A* | 1.018 | 4.4E-02 |
| *METTL7A* | 1.016 | 1.9E-02 |
| *MPP1* | 1.014 | 1.4E-02 |
| *ADGRD1* | 1.014 | 1.4E-02 |
| *COBLL1* | 1.014 | 6.4E-04 |
| *FGF10* | 1.013 | 3.6E-03 |
| *HSD17B14* | 1.013 | 2.3E-02 |
| *SECTM1* | 1.011 | 1.3E-02 |
| *PCSK5* | 1.009 | 4.1E-03 |

| | | |
|---|---|---|
| *SOWAHA* | 1.009 | 3.0E-02 |
| *CHRD* | 1.007 | 2.1E-02 |
| *GATA5* | 1.006 | 3.0E-02 |
| *EDA* | 1.005 | 1.3E-04 |
| *IL2RG* | 1.001 | 9.2E-03 |
| *SGK3* | 1.001 | 4.4E-03 |
